# Csf1, a tunnel-like lipid-transfer protein, mediates lipid remodeling and underpins eukaryotic membrane resilience to high hydrostatic pressure and cold

**DOI:** 10.1101/2025.11.20.689427

**Authors:** Fumiyoshi Abe, Tetsuo Mioka, Yusuke Suzuki, Akane Ogasawara, Daisuke Mochizuki, Saki Imura, Yusuke Kato, Takahiro Mochizuki

## Abstract

Biological membranes continuously remodel their lipid composition to preserve functionality under environmental stress, yet the molecular basis of this process in eukaryotes remains incompletely understood. Here, we identify the tunnel-like lipid transfer protein Csf1 as a central factor mediating adaptive lipid remodeling that enables *Saccharomyces cerevisiae* to tolerate high hydrostatic pressure and low temperature. Quantitative lipidomic and membrane biophysical analyses revealed that loss of Csf1 markedly reduces the unsaturation of phosphatidylserine (PS) and phosphatidylethanolamine (PE), leading to rigidification of the endoplasmic reticulum (ER) membrane. Whereas *OLE1* overexpression partially mitigated this defect at low temperature, no compensatory response occurred under pressure. Overexpression of the PS/phosphatidylinositol 4-phosphate exchanger Osh6/7 restored PS and PE unsaturation and rescued growth of the Csf1-deficient mutant, indicating a cooperative role in sustaining PS flux at ER–plasma membrane (PM) contact sites. Pressure-induced degradation of the amino-acid permease Bap2 further linked lipid imbalance to membrane-protein instability. By supplying unsaturated PS and PE, Csf1 preserves ER and PM flexibility, defining a conserved mechanism of eukaryotic membrane adaptation to extreme physical stress.

## Introduction

Physical stresses such as extreme temperature and hydrostatic pressure alter membrane fluidity, thickness, and phase behavior. To maintain function, organisms remodel their lipid composition through homeoviscous adaptation, which typically involves increasing fatty acid unsaturation at low temperature (Sinensky, 1974; Hazel, 1995; de Mendoza, 2014; Ernst *et al*., 2016; Yu *et al*., 2021). Recent studies of deep-sea invertebrates further introduced the concept of homeocurvature adaptation, in which lipids with negative spontaneous curvature preserve membrane flexibility and fusion under high pressure (Winnikoff *et al*., 2024; Winnikoff and Budin, 2025). Thus, membrane adaptation encompasses both fluidity and curvature control as core strategies for life in extreme environments. Despite this importance, the molecular basis of pressure adaptation remains poorly understood in eukaryotes, whose organelles and membrane-contact–site lipid-transfer systems add complexity to membrane homeostasis.

*Saccharomyces cerevisiae* provides a genetically tractable model for studying how high hydrostatic pressure affects eukaryotic physiology (Abe and Minegishi, 2008; Kurosaka *et al*., 2019; Uemura *et al*., 2020; Abe, 2021; Mochizuki *et al*., 2023; Kato *et al*., 2025). We previously identified Csf1, originally discovered as essential for low-temperature fermentation (Tokai *et al*., 2000), as also critical for growth under high pressure (Abe and Minegishi, 2008). The severe pressure sensitivity of *csf1*Δ cells is rescued when auxotrophic requirements are met (Kurosaka *et al*., 2019), suggesting that Csf1 maintains amino acid uptake under pressure, likely by stabilizing amino acid permeases. Csf1 belongs to the Bridge-like Lipid Transfer Protein (BLTP) superfamily, whose members transfer bulk phospholipids through a conserved hydrophobic groove (Reinisch and Prinz, 2021; Neuman *et al*., 2022; Melia and Reinisch, 2022; Hanna *et al*., 2023). Unlike other BLTPs, Csf1 is anchored to the ER by a single N-terminal transmembrane (TM) helix and functions at multiple organelle contact sites. It supports glycosylphosphatidylinositol anchor synthesis by supplying phosphatidylethanolamine (PE) to the endoplasmic reticulum (ER) (Toulmay *et al*., 2022) and becomes essential when PE transport between mitochondria and peroxisomes is required (John Peter et al., 2022), indicating a broader role in inter-organelle lipid homeostasis. Yet, the physiological directionality of Csf1-mediated lipid flow remains unresolved.

Csf1 homologs are conserved across eukaryotes. In mammals, loss of BLTP1 (Tweek/KIAA1109) causes perinatal lethality with neuromuscular and synaptic defects (Liu *et al*., 2025), and BLTP1 mutations underlie Alkuraya–Kučinskas syndrome, a severe neurodevelopmental disorder (Kumar *et al*., 2020). In *Caenorhabditis elegans*, the homolog LPD-3 forms a giant ER–plasma membrane (PM) bridge that transfers bulk phospholipids and supports cold tolerance; loss of LPD-3 disrupts lipid distribution and represses the fatty-acid desaturase FAT-7, defects rescued by lecithin supplementation (Wang, C. *et al*., 2022). Recent cryo-EM studies show that LPD-3 forms assemblies that accommodate multiple lipids simultaneously, consistent with bulk lipid transfer through its tunnel-like structure (Kang *et al*., 2025). These findings establish Csf1 and its homologs as conserved lipid-transfer factors essential for membrane homeostasis and stress adaptation.

In this study, we used hydrostatic pressure to clarify how Csf1 preserves membrane structure and biophysical homeostasis. Pressure compresses lipid bilayers and increases molecular order, effects that resemble but are mechanistically distinct from those of low temperature. By comparing these two stresses, we show that Csf1 maintains membrane homeostasis via eukaryote-specific lipid-transfer pathways, enabling adaptation to both high pressure and cold. These findings uncover a conserved membrane-remodeling mechanism fundamental to survival in extreme environments.

## Results

### 2.1. Csf1 enables growth under high pressure and low temperature by stabilizing amino acid permeases

In this study, we used a C-terminal truncation mutant (*csf1*ΔC) lacking the region after amino acid 224, generated to avoid affecting transcription of the adjacent essential gene *GAA1* whose promoter overlaps the *CSF1* ORF (Fig. 1A) (John Peter *et al*., 2022). The *csf1*ΔC strain grew normally under the control condition (0.1 megapascals [MPa], 1 bar, 0.9869 atm, 1.0197 kg/cm^2^; to avoid confusion, we have consistently used MPa throughout) but showed pronounced growth defects under high pressure (25 MPa, 25°C) and low temperature (0.1 MPa, 15°C). These results demonstrate that the stress sensitivity reflects loss of Csf1 itself. Deletion of genes encoding other BLTP-related proteins (Vps13, Atg2, Fmp27, and Hob2) did not impair growth under high pressure or low temperature, indicating a distinct and nonredundant role for Csf1 (Fig. 1B). Similar to the previously reported deletion mutants of the poorly characterized genes *EHG1/MAY24, MTC2, MTC4, MTC6,* and *DLT1* (Kurosaka *et al*., 2019), the *csf1*ΔC strain regained high-pressure and cold tolerance when auxotrophic requirements were met by introduction of the pHLUK plasmid (Fig. 1B). These findings suggest that the stress sensitivity of *csf1*ΔC cells results primarily from the loss or destabilization of nutrient permeases.

**Fig. 1.**
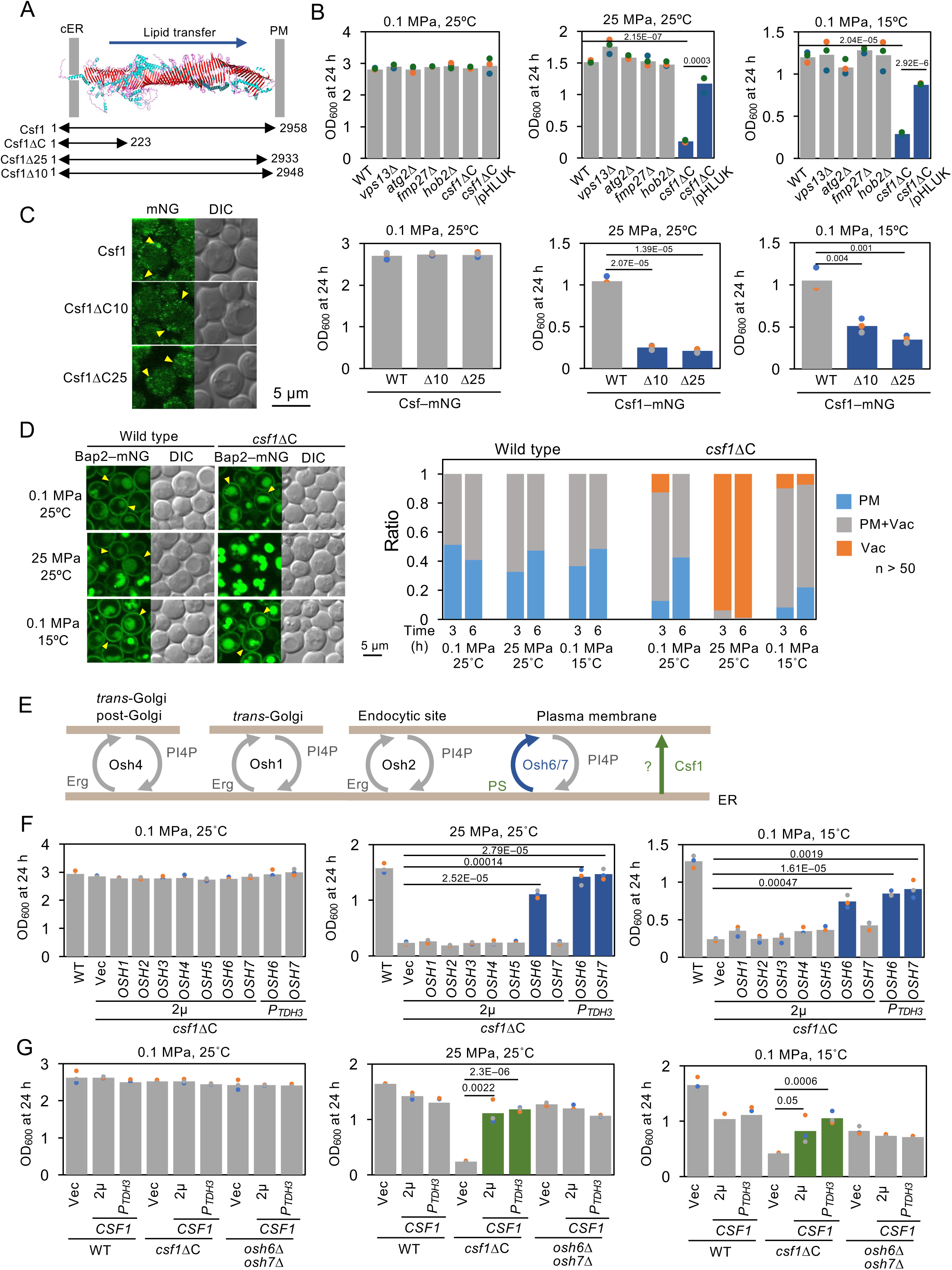
Csf1-dependent growth under high pressure and low temperature, and rescue of the *csf1*ΔC defect by *OSH6/OSH7* overexpression. (A) Predicted three-dimensional structure of Csf1 generated using AlphaFold2 (Jumper et al, 2021; Varadi et al, 2022) and schematic representation of the deletion constructs used in this study. (B) Growth defects of the *csf1*ΔC strain compared with the wild-type (WT) strain and other BLTP mutants (*vps13*Δ, *fmp27*Δ, *hob2*Δ, *atg2*Δ,) under the indicated conditions, assessed by OD_600_ after 24 h. Complementation with pHLUK (*HIS3 LEU2 URA3 LYS2*) restored the growth of the *csf1*ΔC mutant under high-pressure or low-temperature conditions. Strains with statistically significant differences are highlighted in blue. (C) Comparable localization of Csf1Δ10–mNG and Csf1Δ25–mNG to WT Csf1–mNG (yellow arrowheads). Growth defects of *csf1*Δ10 and *csf1*Δ25 mutants compared with WT cells. Data are shown as mean values (bars) with individual OD_600_ points from three independent experiments. Strains with statistically significant differences are highlighted in blue. (D) Loss of Csf1 destabilizes Bap2 at the PM. Subcellular localization of Bap2–mNG in wild-type and *csf1*ΔC cells after 3 h of culture under the indicated conditions. Yellow arrowheads indicate Bap2 at the PM (left). Quantification of Bap2 localization among the PM (PM), PM + vacuole (Vac), and vacuole (Vac) alone after 3 h and 6 h of incubation (right). In *csf1*ΔC cells, Bap2 predominantly shifted to the vacuole, especially after high-pressure culture (n > 50 cells). (E) Schematic illustration of the intracellular localization and lipid-transfer activity of Osh proteins. (F) Growth of *csf1*ΔC cells overexpressing *OSH6* or *OSH7* from a multicopy (2µ) plasmid or under the *TDH3* promoter. Strains with statistically significant differences are highlighted in blue. (G) Growth of *osh6*Δ*osh7*Δ cells overexpressing *CSF1* from a multicopy plasmid or under the *TDH3* promoter. Data are shown as mean values (bars) with individual OD_600_ points from three independent experiments. Strains with statistically significant differences are highlighted in green.

Csf1–mNeonGreen (mNG) localized to peripheral puncta that partially overlapped with the cortical ER (cER) marker Rtn1–mCherry (Fig. EV1). Truncation mutants lacking only 10 or 25 C-terminal residues localized to the cER, yet even the 10-residue deletion caused strong high-pressure and cold sensitivity (Fig. 1C). These results indicate that physical contact between the C-terminal region and the PM is essential for the lipid-transfer activity of Csf1.

To determine whether loss of Csf1 affects permease stability, we examined the localization of the branched-chain amino acid permease Bap2–mNG. Under the control condition, Bap2 localized to the PM and vacuole in both strains, consistent with normal turnover. Under high pressure, Bap2 rapidly disappeared from the PM and accumulated in the vacuole within 3 h in *csf1*ΔC cells (Fig. 1D). These observations show that Csf1 supports growth under high pressure by stabilizing amino acid permeases at the PM, likely by maintaining an appropriate membrane lipid environment. However, such degradation was not evident at low temperature, consistent with lipidomic data described later.

### 2.2. Overexpression of *OSH6* and *OSH7* suppresses the high-pressure and low-temperature sensitivity of the *csf1*ΔC mutant

We examined the oxysterol-binding protein-related proteins Osh1 to Osh7, which mediate lipid exchange at membrane contact sites (Stefan *et al*., 2011, Reinisch and Prinz, 2021)(Fig. 1E). Overexpression of *OSH6* from a multicopy plasmid markedly restored the growth of the *csf1*ΔC strain at high pressure and low temperature, whereas *OSH1* to *OSH5* and *OSH7* had little effect (Fig. 1F). However, strong *OSH7* expression from the *TDH3* promoter produced a similar rescue. Osh6 and Osh7 mediate phosphatidylserine (PS)/phosphatidylinositol-4-phosphate (PI4P) exchange at cER–PM contact sites (Moser von Filseck *et al*., 2015), suggesting a possible functional connection to Csf1. However, genetic analyses revealed a clear asymmetry between these systems: *OSH6* or *OSH7* overexpression suppressed the high-pressure sensitivity of *csf1*ΔC, but the mild sensitivity of the *osh6*Δ*osh7*Δ double mutant was not rescued by *CSF1* overexpression (Fig. 1G). These results indicate that Csf1 does not directly catalyze PS/PI4P exchange like Osh6 and Osh7. Instead, they function within a shared lipid-transfer network, but with nonequivalent roles. Under high-pressure and low-temperature conditions, Csf1 likely acts upstream to remodel membrane lipid composition and optimize membrane physical properties, thereby enabling Osh6 and Osh7 to maintain efficient PS/PI4P exchange and membrane homeostasis.

### 2.3 Defective lipid remodeling and maintenance of membrane curvature in the *csf1*ΔC mutant

We analyzed lipid composition in wild-type, *csf1*ΔC, and *csf1*ΔC/*P_TDH3_*-*OSH6* (hereafter *csf1*ΔC/*OSH6*) strains cultured for 16 h under control (Ctrl, 0.1 MPa, 25 °C), high-pressure (HP, 25 MPa, 25 °C), and low-temperature (LT, 0.1 MPa, 15 °C) conditions. According to the criteria described in the Materials and methods section, we used the P100 fraction as an ER-enriched microsomal fraction. Western blotting confirmed that the distributions of Pma1 (PM marker), Dpm1 (ER marker), and Pep12 (endosomal marker) were similar among strains, indicating comparable membrane recovery (Fig. EV2).

Principal-component analysis (PCA) showed that under high pressure, the *csf1*ΔC samples formed a distinct cluster compared with all other strains and conditions, indicating that loss of Csf1 exerts the strongest impact specifically at high pressure (Fig. 2A). Lipid composition was markedly altered in the *csf1*ΔC mutant under both high pressure and low temperature, and these changes were partially alleviated by *OSH6* overexpression (Fig. EV3A; Tables EV1 and EV2). To interpret the physical implications of these complex compositional shifts, we applied the “homeocurvature” framework proposed by Winnikoff et al. (2024) (Fig. 2B). In this model, the phospholipid curvature index (PLCI) is calculated as the weighted average of the intrinsic curvature (*c*_0_) values of each lipid species multiplied by their abundance (Winnikoff *et al*., 2024; Winnikoff and Budin, 2025)(see Materials and methods). Under the control condition, PLCI did not differ among the three strains (Fig. 2B). After high-pressure cultivation, PLCI increased in wild-type cells, suggesting membrane flattening, but decreased in the *csf1*ΔC strain, indicating abnormally enhanced negative curvature (Fig. 2B). *OSH6* overexpression did not correct this defect. At low temperature, PLCI was higher in *csf1*ΔC than in wild-type cells, showing the opposite trend (Fig. 2B). In deep-sea ctenophores, lower PLCI reflects enrichment of lipids with negative curvature that promote pressure tolerance (Winnikoff *et al*., 2024), whereas yeasts exhibited pressure-induced membrane flattening that was lost in the *csf1*ΔC mutant. These results suggest that in yeasts, membrane-shaping proteins such as reticulons contribute more to curvature control than lipid composition alone (West *et al*., 2011; Wang, N. *et al*., 2021). Taken together, these findings demonstrate that Csf1 is required for coordinated lipid remodeling and curvature maintenance, allowing cellular membranes to maintain their optimal physical state under high pressure and low temperature.

**Fig. 2.**
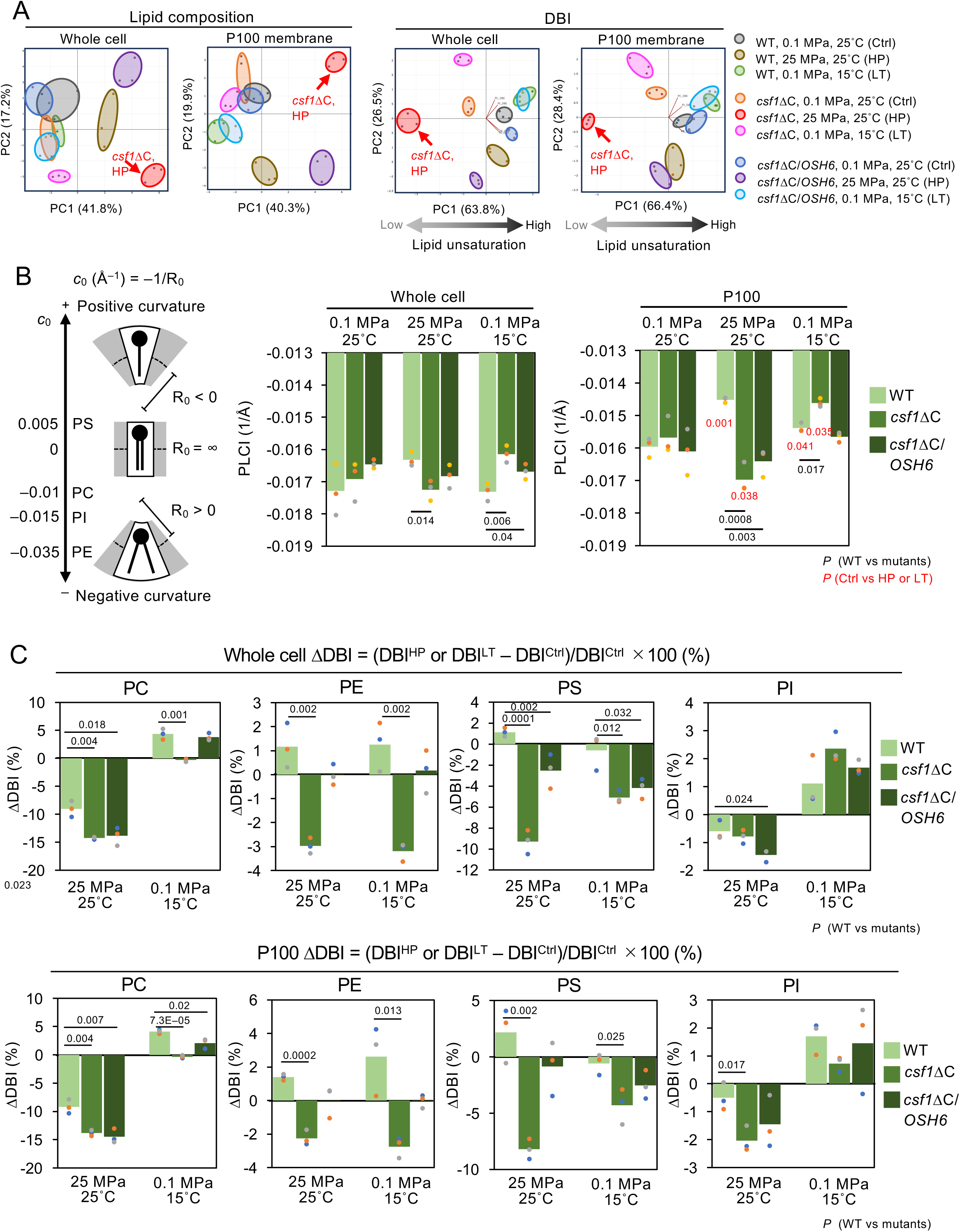
Csf1 preserves phospholipid composition and unsaturation under high pressure and low temperature. (A) Principal component analysis (PCA) of major phospholipids (PC, PE, PI, PS) and their double-bond index (DBI) in whole-cell and P100 membrane fractions. In both analyses, the *csf1*ΔC mutant (red arrows) cultured under high pressure shows a marked divergence from the other samples. PCA was performed using the Statistics Kingdom web tool. (B) Loss of Csf1 exerts complex and condition-dependent effects on membrane curvature under high pressure and low temperature. (Left) Intrinsic curvature (*c*_0_) is a measure of lipid molecular shape, defined as the reciprocal of the radius of a mechanically unstressed monolayer (*R*_0_) at its neutral plane of bending (dashed line). The illustration was adapted from (Winnikoff and Budin, 2025) with modifications. The reference *c*_0_ values were adopted from (Dymond, 2021) and (Mulet *et al*., 2008)(PC = –0.01, PE = –0.035, PS = 0.005, PI = –0.015; Å^−1^). (Right) Estimation of membrane curvature. The phospholipid curvature index (PLCI) was calculated for each strain under the indicated conditions, based on the reference *c*_0_ values. (C) DBI values of major phospholipids in whole-cell and P100 membrane fractions. Loss of Csf1 reduced DBI, particularly in PS and PE after high-pressure culture, whereas *OSH6* overexpression restored unsaturation levels. Data are shown as mean values (bars) with individual points from three independent preparations. Absolute values of the bar graphs are provided in Table EV3 and EV4.

### 2.4 Abnormal lipid-desaturation response in the *csf1*ΔC mutant and its recovery by *OSH6* overexpression

To assess how Csf1 deficiency affects membrane unsaturation, we quantified the double-bond index (DBI) for each phospholipid class. PCA based on DBI values showed that *csf1*ΔC cells cultivated under high pressure formed a cluster distinct from those of the other two strains and conditions (Fig. 2A, Fig. EV3B and in Tables EV3 and EV4). To evaluate desaturation changes, we calculated the difference in DBI (ΔDBI) between the control condition and either high pressure or low temperature (Fig. 2C).

Under high pressure, wild-type cells exhibited a substantial DBI decrease in phosphatidylcholine (PC) (−9%) in whole-cell lipids, whereas PE, PS, and PI remained essentially unchanged (~1%), indicating that PC primarily buffers membrane fluidity (Fig. 2C). In contrast, the *csf1*ΔC mutant exhibited a global decline in whole-cell lipid unsaturation: PC decreased by 14%, all other classes were reduced, and PS exhibited a substantial drop (−9%), revealing a collapse of normal remodeling (Fig. 2C). *OSH6* overexpression markedly restored unsaturation in PS and PE, consistent with Osh6-mediated PS/PI4P exchange at cER–PM contact sites (Fig, 2C). Although Osh6 does not transport PE, the recovery of PE unsaturation likely reflects indirect improvement of the membrane environment through restored PS flux.

Under low temperature, the pattern differed. In wild-type cells, the DBI of PC increased by about 8%, consistent with activation of the fatty acid desaturase Ole1. In *csf1*ΔC cells, the increase was minimal (0 to 2%), suggesting that abnormal ER organization interferes with Ole1 mediated desaturation. *OSH6* overexpression enhanced the DBI of PC (Fig. 2C). For PE, *csf1*ΔC showed a 3% decrease, similar to the change observed under high pressure, whereas PS decreased only moderately. *OSH6* overexpression restored PE unsaturation but had little effect on PS. The trends in whole cell lipidomes were largely mirrored in the P100 membrane fraction (Fig. 2C).

These results indicate that Csf1 maintains a pool of highly unsaturated lipids under both stress conditions. Notably, in both strains, the direction of PC unsaturation differed between stresses. At low temperature, PC unsaturation was preserved even without Csf1, which may explain the stability of Bap2 at the PM (Fig. 1D). The rescue of low temperature growth by pHLUK (Fig. 1B) therefore likely reflects metabolic compensation of substrate transport rather than suppression of permease degradation.

To test whether the DBI decrease in *csf1*ΔC results from impaired fatty acid desaturation, we examined the desaturase Ole1 (Stukey et al., 1990). In both strains, Ole1–mNG localized to the ER and its fluorescence decreased comparably after 6 h at high pressure (Fig. 3A), indicating that reduced DBI in *csf1*ΔC is not due to diminished Ole1 expression. After low-temperature cultivation, Ole1 fluorescence increased in both strains, showing that the Mga2/Spt23 regulatory pathway (Covino et al., 2016; Ballweg et al., 2020) remains functional (Fig. 3A). Overexpression of *OLE1* produced only a negligible improvement in *csf1*ΔC growth under high pressure (approximately 20% of the wild-type level) but substantially enhanced growth at low temperature (approximately 62%) (Fig. 3B). Thus, wild-type cells maintain lipid unsaturation under pressure without upregulating Ole1, whereas *csf1*ΔC cells cannot. Csf1 therefore likely supports a remodeling process independent of Ole1 during pressure adaptation, whereas at low temperature, *OLE1* overexpression compensates through a Csf1-independent pathway (Fig. 3B).

**Fig. 3.**
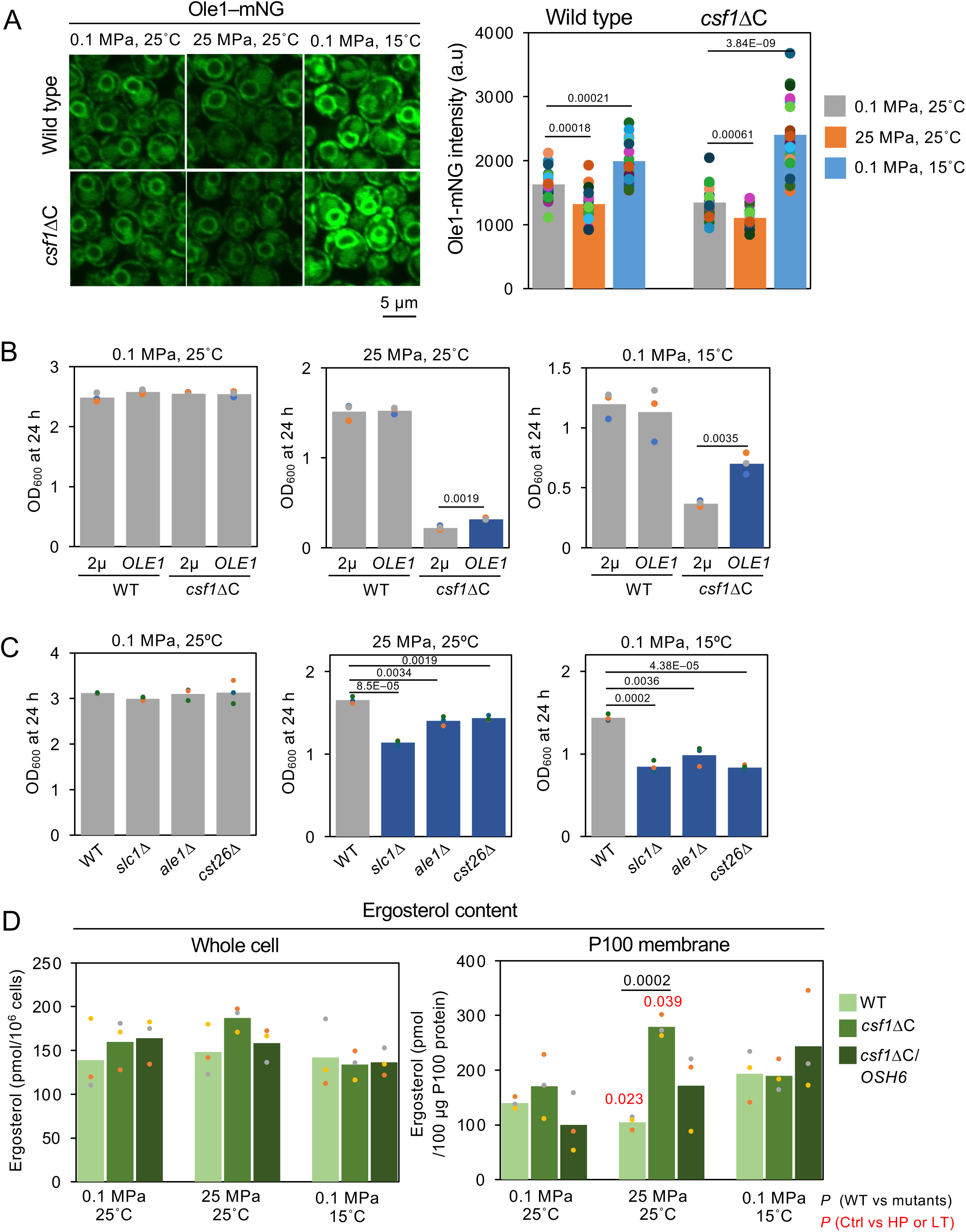
Distinct effects of *OLE1* overexpression and membrane ergosterol content on adaptation to high pressure and low temperature. (A) Ole1–mNG fluorescence intensity in wild type and *csf1*ΔC cells after 6 h of culture under the indicated conditions. Ole1 expression increased at low temperature but decreased under high pressure in both strains (n = 20 cells for each condition). (B) Overexpression of *OLE1* in *csf1*ΔC cells restored growth under low temperature but not under high pressure. Data are shown as mean values (bars) with individual OD_600_ points from three independent experiments. Strains with statistically significant differences are highlighted in green. (C) Deletion mutants of phospholipid acyltransferase genes show moderate high-pressure or low-temperature sensitivity. Data are shown as mean values (bars) with individual points from three independent preparations. Strains with statistically significant differences are highlighted in green. (D) Lipidomic analysis of ergosterol content using the same samples as those analyzed for phospholipid. The *csf1*ΔC strain cultured under high pressure exhibited a marked increase in ergosterol levels in the P100 membrane fraction.

To place Csf1 function in the context of lipid-remodeling pathways, we analyzed the *slc1*Δ, *ale1*Δ, and *cst26*Δ mutants under the same stress conditions. These mutants showed mild growth defects distinct from the severe sensitivity of *csf1*ΔC (Fig. 3C). Slc1, Ale1, and Cst26 function in the Lands’ cycle reacylation of lysophospholipids (Patton-Vogt and de Kroon, 2020). Their modest phenotypes suggest that although the Lands’ cycle contributes to membrane maintenance, it is insufficient for full adaptation to pressure and cold. Csf1 likely cooperates with this pathway to regulate broader lipid fluxes and stabilize membrane architecture under extreme stress.

Under high pressure, *csf1*ΔC cells showed increased lipid saturation, suggesting higher membrane order. Because such ordering can influence sterol behavior, we quantified ergosterol, a key determinant of membrane rigidity. In whole-cell extracts, no significant changes occurred after pressure or cold exposure (Fig. 3D). However, in the P100 fraction, *csf1*ΔC cells accumulated approximately 1.6-fold more ergosterol after high-pressure cultivation, and this increase was reversed by *OSH6* overexpression (Fig. 3D). At low temperature, ergosterol levels remained unchanged. These findings suggest that in *csf1*ΔC cells, reduced lipid unsaturation, particularly enrichment of PE and loss of PS unsaturation in the ER, leads to membrane rigidification that promotes local ergosterol accumulation.

### 2.5 Loss of Csf1-dependent lipid unsaturation decreases ER membrane flexibility, whereas OSH6 overexpression compensates for this defect

To assess how the compositional change affects membrane physical properties, we quantitatively analyzed membrane order and fluidity by time-resolved fluorescence anisotropy using trimethylammonium-diphenylhexatriene (TMA-DPH)(Fig. 4A)(Abe and Hiraki, 2009; Abe *et al*., 2009). In living cells, the probe partitions exclusively into the outer leaflet of the PM, allowing selective evaluation of PM order (Sheridan and Block, 1988; Abe and Hiraki, 2009).

**Fig. 4.**
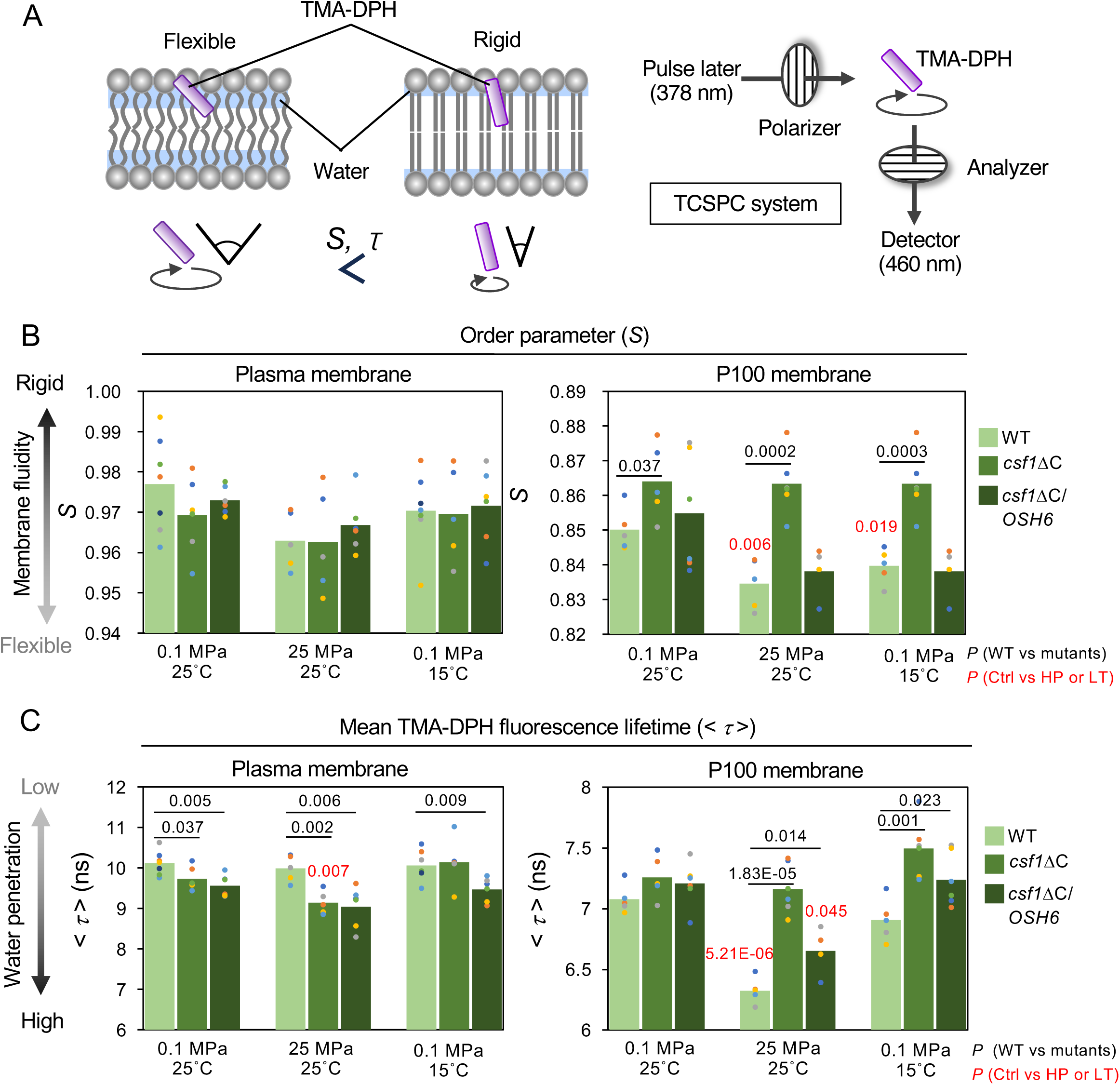
Csf1 is required for membrane flexibility as assessed by TMA–DPH fluorescence anisotropy measurements. (A) Schematic representation of the time-resolved TMA–DPH fluorescence anisotropy measurements based on time-correlated single-photon counting (TCSPC). (B) Order parameter *S* and (C) mean fluorescence lifetime of TMA–DPH (< τ >) measured in whole cells and P100 membranes. Cells were cultured as in Fig. 3. Measurements were performed under atmospheric pressure. Wild-type P100 membranes became more flexible after growth under high pressure or low temperature, whereas P100 membranes of *csf1*ΔC cells remained rigid under the same conditions. Overexpression of *OSH6* under the control of the *TDH3* promoter restored membrane flexibility in *csf1*ΔC cells. Data are shown as mean values (bars) with individual values from more than four independent experiments. TMA–DPH, trimethylammonium-diphenylhexatriene.

Measurements showed that *S* values did not differ significantly between wild-type and *csf1*ΔC cells under any condition (Fig. 4B), suggesting that alterations in phospholipid composition were not detectable at the bulk PM level. This may be because the high ergosterol content of the PM dominates membrane ordering. In contrast, pronounced differences were observed in the P100 fraction representing the ER membrane. P100 membranes of wild-type cells displayed lower *S* values than the PM, consistent with their inherently flexible nature. Cultivation under high pressure or low temperature further decreased *S* in the wild-type strain, indicating activation of homeoviscous adaptation that restores ER flexibility under stress (Fig. 4B). However, the *csf1*ΔC mutant exhibited persistently high *S* values under all conditions, showing complete loss of this membrane-softening response. Remarkably, *OSH6* overexpression restored *S* in *csf1*ΔC P100 membranes to wild-type levels, demonstrating that enhanced lipid transport at ER–PM contact sites compensates for the flexibility defect caused by Csf1 loss (Fig. 4B).

The <*τ*> values paralleled these findings. In wild-type cells, cultivation under high pressure decreased <*τ*>, reflecting greater water penetration into the headgroup region associated with membrane softening. In *csf1*ΔC cells, <*τ*> was higher than that in wild-type cells, indicating exclusion of water due to membrane rigidification (Fig. 4C). In *csf1*ΔC/*OSH6* cells, <*τ*> partially recovered after cultivation under high pressure or low temperature (Fig. 4C). Together, these findings show that loss of Csf1 disrupts ER membrane flexibility, whereas *OSH6* overexpression compensates for this defect through enhanced lipid exchange at ER–PM contact sites (Fig. 5).

**Fig. 5.**
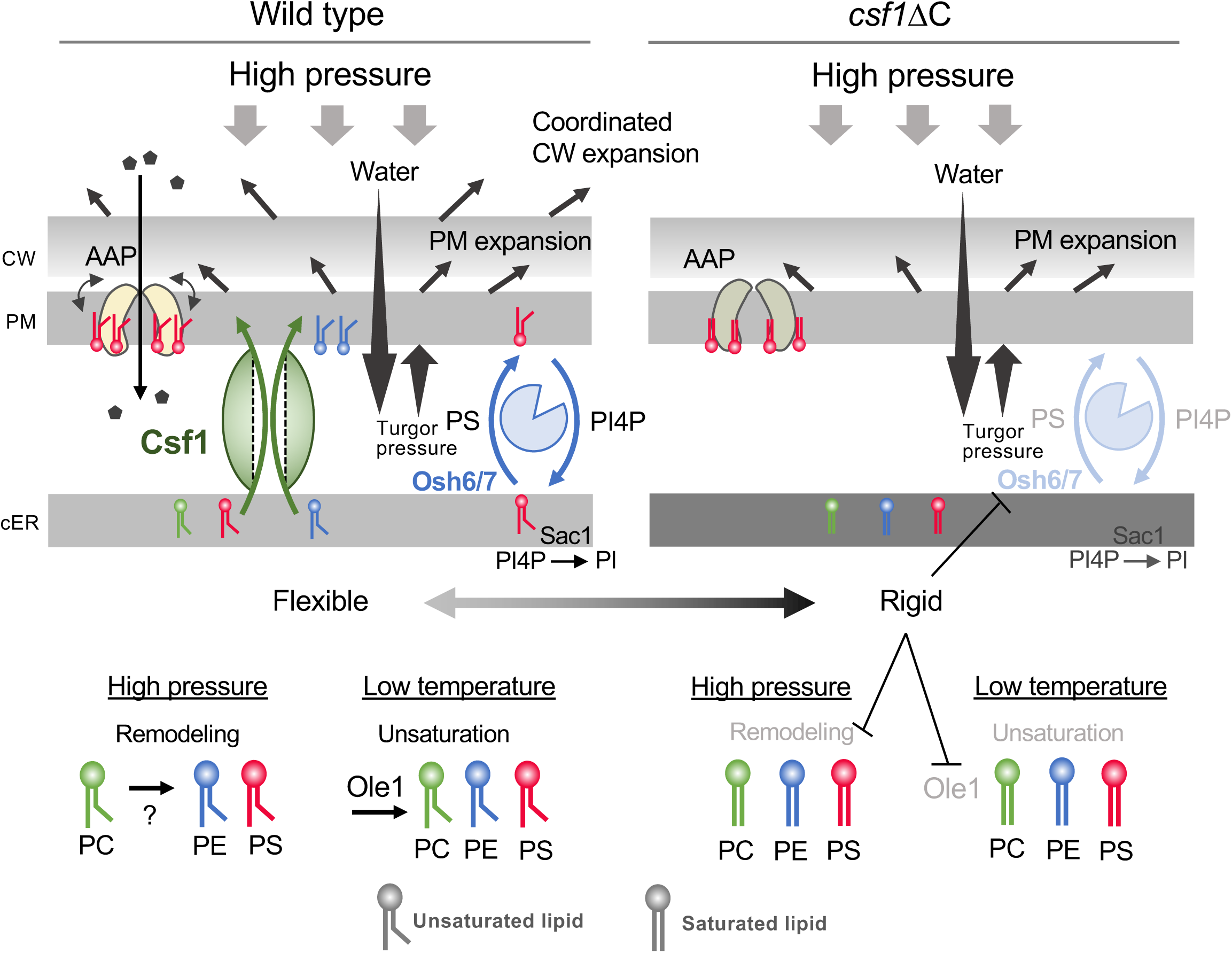
Working model of Csf1-dependent membrane adaptation to high pressure and low temperature. Left panel: In wild-type cells, the ER maintains high lipid unsaturation. At ER–PM contact sites, the tunnel-like lipid-transfer protein Csf1 supplies unsaturated PS and PE from the ER to the PM, while Osh6/7 drive PS/PI4P exchange through Sac1-dependent PI4P turnover. These coordinated fluxes maintain a continuous supply of unsaturated lipids, support ER flexibility (low *S* value), and stabilize amino acid permeases through PS-rich boundary lipid shells. Under high pressure, accelerated lipid remodeling and pressure-driven PM stretching may facilitate Csf1-mediated bulk transfer. At low temperature, Ole1 induction helps preserve PS and PE unsaturation. Right panel: In the csf1ΔC mutant, the Csf1-dependent bulk lipid supply is lost. Under high pressure or cold, tighter ER packing raises the energetic barrier for PS extraction, compromising Osh6/7-mediated PS/PI4P exchange. As a result, the DBI of PS, PE, and PC declines, the ER becomes rigid (high *S* value), and permeases that require unsaturated PS for structural flexibility are destabilized and degraded in the vacuole. Overexpression of *OSH6* partially restores PS and PE unsaturation, improves ER flexibility, and rescues the growth defect. These observations indicate that Csf1 maintains ER unsaturation through ER–PM lipid flux and acts upstream of Osh6/7 in a hierarchical pathway that ensures adaptive remodeling at the ER–PM interface under high-pressure and low-temperature stress.

## 3. Discussion

This study demonstrates that Csf1 is essential for membrane remodeling in *S. cerevisiae* under high pressure and low temperature. High hydrostatic pressure provides a uniquely sensitive condition that reveals the vulnerability of the *csf1*ΔC mutant and clarifies the physiological role of Csf1 (Fig. 5). At low temperature, reduced lipid unsaturation in *csf1*ΔC cells is partially offset by *OLE1* induction through the Mga2/Spt23 pathway (Covino *et al*., 2016; Ballweg *et al*., 2020), masking Csf1 dependence. In contrast, high pressure suppresses *OLE1* expression and removes this compensation (Fig. 3). Thus, pressure sensitivity offers a clear readout of Csf1 function, linking lipid remodeling to membrane-protein stability. High hydrostatic pressure compresses the free volume of lipid bilayers, reduces gauche conformations, increases molecular order and membrane thickness, and decreases the area per lipid molecule (Winter and Jeworrek, 2009). Low temperature also increases membrane order, but through reduced thermal motion rather than volumetric compression, making the two effects mechanistically distinct. Given that the Mga2 sensor detects local packing changes around its transmembrane residue Trp1042 rather than global fluidity (Ballweg *et al*., 2020), localized ordering at low temperature is likely sensed, whereas uniform ordering under high pressure diminishes structural fluctuations and weakens sensor activation.

Lipidomic and biophysical analyses showed that wild-type cells maintain selective remodeling under pressure, decreasing unsaturation of PC while preserving that of PS, PE, and PI (Fig. 2). In *csf1*ΔC cells, unsaturation decreased across all classes, with marked reductions in PS and PE, and the ER membrane became rigid. PC likely acts as a reservoir for unsaturated acyl chains, replenishing PS and PE via the Lands’ cycle (Patton-Vogt and de Kroon, 2020). In tightly packed membranes, reduced extractability of PS or PI4P would impair Osh6/7 mediated exchange, making Csf1 dependent bulk lipid transfer critical for sustaining lipid flux and membrane homeostasis (Fig. 5). Our findings align with previous reports that Csf1 localizes to cER–PM contact sites and is required for cold adaptation (John Peter *et al*., 2022), and we extend this role to high pressure, underscoring a conserved requirement for maintaining ER–PM lipid flux.

Csf1 is unlikely to transport a single lipid species; rather, it may facilitate PS and PE delivery to the PM as part of broader ER compositional shifts. Although the driving force remains unclear, our previous report suggests a plausible biophysical basis. Exposure to 25 MPa causes a ~1.2-fold increase in cell volume due to water influx, elevating turgor pressure and stretching the PM (Mochizuki *et al*., 2023). This stretching reduces local packing density and may generate a lipid concentration gradient between the cER and PM (cER > PM), which Csf1 could exploit to draw lipids into its tunnel-like interior. As cells grow, this mechanically coupled process may coordinate lipid transfer with membrane expansion and cell-wall elongation, promoting adaptation under high pressure (Fig. 5).

Periprotein lipidomic analyses show that amino acid permeases reside in membranes that are depleted of ergosterol but enriched in unsaturated phospholipids, especially PS (van ’t Klooster *et al*., 2020a; van ’t Klooster *et al*., 2020b). This anionic lipid shell provides the flexibility required for the conformational changes associated with substrate transport. Therefore, the Csf1 dependent supply of unsaturated PS is directly linked to permease stability, which becomes particularly important under high pressure and low temperature when membrane packing is enhanced (Fig. 5).

Recent studies propose a cooperative “bridge and scramblase” model in which BLTPs and lipid scramblases maintain lipid flow and bilayer equilibrium (Melia and Reinisch, 2022; Hanna *et al*., 2023). Structural modeling (Fig. EV4) and our proteomic analyses (unpublished observation) indicate that Csf1 associates with Ehg1/May24, Mtc2, and Dlt1, and deletion of any of these genes causes similar high-pressure and low-temperature sensitivity (Abe and Minegishi, 2008; Kurosaka *et al*., 2019), suggesting that they operate as a shared module. Ehg1/May24 also stabilizes the tryptophan permease Tat2 under high pressure (Kato *et al*., 2025). Consistent with this view, Rocha-Roa et al. (2025) showed physical interactions among Csf1, Ehg1, and Dlt1, and molecular dynamics simulations demonstrated that Ehg1 has lipid-scrambling activity comparable to the human ER scramblase TMEM41B (Rocha-Roa *et al*. 2025). Together, these findings support a conserved Csf1–Ehg1/May24 module that integrates bulk lipid transfer with bilayer re-equilibration. We did not attempt to biochemically reconstitute Csf1-mediated lipid transfer in this study, because our primary aim was to clarify how Csf1-dependent lipid remodeling affects ER and PM biophysical properties under high pressure and cold in living cells. We are currently establishing an in vitro membrane-reconstitution system to directly assess the lipid-transfer activity of Csf1 and its cooperation with Osh6/7 and ER-resident scramblases, and these mechanistic analyses will be presented in a separate study. While BLTP-mediated transport can be bidirectional, the metabolic and chemical gradients of PS and PE favor movement from the cortical ER to the PM. We therefore infer that under high-pressure stress, Csf1 primarily supplies unsaturated PS and PE to the PM. Together, these considerations support a model in which high-pressure stress increases the cellular demand for unsaturated PS and PE at the PM, making Csf1-mediated lipid supply essential (Fig. 5).

In summary, this study proposes that Csf1 cooperates with Osh6/7 to sustain lipid exchange between the ER and PM, thereby preserving membrane flexibility and amino acid permease function. This model links global homeoviscous adaptation with local control of the permease microenvironment and explains how eukaryotic membranes remain resilient under extreme pressure and cold. Our findings also connect environmental adaptation to human diseases caused by defects in membrane homeostasis. Csf1 thus emerges as a unifying factor that integrates the physical principles of membrane adaptation with the pathophysiology of lipid-related disorders.

## 4. Materials and methods

### 4.1. Yeast strains and culture conditions

The yeast strains used in this study are listed in Table 1. Strain BY4742 (*MAT***α** *his3*Δ*1 leu2*Δ*0 lys2*Δ*0 ura3*Δ*0*) was used as a parental wild-type strain. Cells were grown at 25 °C in synthetic complete medium consisting of 0.67% yeast nitrogen base without amino acids, supplemented with adenine sulfate (20 µg/mL), uracil (20 µg/mL), tryptophan (40 µg/mL), histidine-HCl (20 µg/mL), leucine (90 µg/mL), lysine-HCl (30 µg/mL), arginine-HCl (20 µg/mL), tyrosine (30 µg/mL), isoleucine (30 µg/mL), phenylalanine (50 µg/mL), glutamic acid (100 µg/mL), aspartic acid (100 µg/mL), threonine (200 µg/mL), serine (400 µg/mL), and 2% D-glucose, with additional nutrients added as required to complement auxotrophies. For cultivation under high hydrostatic pressure or low temperature, exponentially growing cultures were diluted in synthetic complete medium to an OD_600_ of 0.1. The diluted cultures were dispensed into sterilized 1.7 mL tubes, sealed with Parafilm, and subjected to high pressure (25 MPa, 25 °C) in hydrostatic chambers (Abe and Minegishi, 2008) or incubated at 0.1 MPa and 15 °C. After incubation, the pressure was released, and cell growth was assessed by measuring the OD_600_ using a PD-307 spectrophotometer (Apel, Kawaguchi, Japan). An OD_600_ of 0.1 corresponds to 1.65 × 10^6^ cells/mL.

**Table 1.** Strains used in this study.

| Strains | Genotype | Source or reference |
| --- | --- | --- |
| BY4742 | *MATa his3Δ1 leu2Δ0 lys2Δ0 ura3Δ0* | (Brachmann *et al.*, 1998) |
| *csf1ΔC* | *csf1ΔC::kanMX4* in BY4742 | This study |
| *vps13Δ* | *vps13Δ::kanMX4* in BY4742 | (Winzeler *et al.*, 1999) |
| *atg2Δ* | *atg2Δ::kanMX4* in BY4742 | (Winzeler *et al.*, 1999) |
| *fmp27Δ* | *fpm27Δ::kanMX4* in BY4742 | (Winzeler *et al.*, 1999) |
| *hob2Δ* | *hob2Δ::kanMX4* in BY4742 | (Winzeler *et al.*, 1999) |
| *osh6Δosh7Δ* | *osh6Δ::kanMX4 osh7Δ::hphMX6* in BY4742 | This study |
| *slc1Δ* | *slc1ΔC::kanMX4* in BY4742 | (Winzeler *et al.*, 1999) |
| *ale1Δ* | *ale1ΔC::kanMX4* in BY4742 | (Winzeler *et al.*, 1999) |
| *cst26Δ* | *cst26ΔC::kanMX4* in BY4742 | (Winzeler *et al.*, 1999) |
| Csf1–mNG | *CSF1–mNG::hphMX6* in BY4742 | This study |
| Csf1Δ10–mNG | *CSF1Δ10–mNG::hphMX6* in BY4742 | This study |
| Csf1Δ25–mNG | *CSF1Δ25–mNG::hphMX6* in BY4742 | This study |
| Csf1–mNG, Rtn1– | *CSF1–mNG::hphMX6, RTN1–* |  |
| mCherry | *mCherry::URA3* in BY4742 | This study |
| BY4742/Ole1– |  |  |
| mNG | *OLE1–mNG::hphMX6* in BY4742 | This study |
| *csf1ΔC*/Ole1–mNG | *OLE1–mNG::hphMX6* in *csf1ΔC* | This study |

### 4.2. Construction of plasmids and strains

Plasmids were constructed according to standard protocols and are listed in Table 2. Using genomic DNA from strain BY4742 as a template, the open reading frames and promoters of *CSF1*, *OSH1–7,* and *OLE1* were amplified by polymerase chain reaction (PCR) and cloned into YEplac195 (*URA3* 2µ). The open reading frames of *CSF1*, *OSH6*, and *OSH7* were also cloned into pUA35 (*TDH3* promoter, *URA3 CEN*). PCR-based gene disruption and gene tagging were performed essentially as described previously (Longtine *et al*., 1998). The essential gene *GAA1* is encoded on the complementary strand upstream of *CSF1*, with its start codon located only 376 bp away from that of *CSF1*. To avoid interference with GAA transcription, a 1-kb upstream region including the *GAA1* promoter was preserved, and disruption of *CSF1* was initiated 670 bp downstream of its start codon. This was achieved using primers CSF1.C2649aaDel-F1-a (5’– GAAACAGATATCGGCAAACATCCAAAGCTACTGATGTTTTcggatccccgggttaattaa–3’) and CSF1-Ctag-b (5’– ACATAAACCAGAAATATGGTATCAAGGACTTTTGAATATAATTAGGAACGgaattcgagctcgtttaaac–3’), together with the template plasmid pFA6a-kanMX6, generating the *csf1*ΔC strain. C-terminally truncated Csf1 variants fused to–mNG were also constructed. A strain lacking the last 10 amino acids (Csf1Δ10–mNG) was generated using primers CSF1-10del-Ctag-a (5’–AAACCGCAGTTATCCCCGTCCAAAAGCTTGTTTATCTTGCAGAAAAGCAGcggatccccgggttaatt aa–3’) and CSF1-Ctag-b. Similarly, a strain lacking the last 25 amino acids (Csf1Δ25–mNG) was obtained using primers CSF1-25del-Ctag-a (5’–AATGGTTTGGTGTCAATAGAAAAAAATTTCCGAAATTCACTCACCAAACCcggatccccgggttaatt aa–3’) and CSF1-Ctag-b.

**Table 2.** Plasmids used in this study.

| Plasmid | Description | Source or reference |
| --- | --- | --- |
| pHLUK | *HIS3 LEU2 URA3 LYS2 CEN6* | addgene |
| pUA35 | *TDH3* promoter, *URA3 CEN4* | Laboratory stock |
| YEplac195 | *URA3* 2μ | (Gietz and Sugino, 1988) |
| pRS316 | *URA3 CEN6* | (Sikorski and Hieter, 1989) |
| YEplac195–CSF1 | *CSF1* driven by its own promoter, *URA3* 2μ | This study |
| pUA35–CSF1 | *CSF1* driven by the *TDH3* promoter, *URA3* 2μ | This study |
| YEplac195–OSH1 | *OSH1* driven by its own promoter, *URA3* 2μ | This study |
| YEplac195–OSH2 | *OSH2* driven by its own promoter, *URA3* 2μ | This study |
| YEplac195–OSH3 | *OSH3* driven by its own promoter, *URA3* 2μ | This study |
| YEplac195–OSH4 | *OSH4* driven by its own promoter, *URA3* 2μ | This study |
| YEplac195–OSH5 | *OSH5* driven by its own promoter, *URA3* 2μ | This study |
| YEplac195–OSH6 | *OSH6* driven by its own promoter, *URA3* 2μ | This study |
| YEplac195–OSH7 | *OSH7* driven by its own promoter, *URA3* 2μ | This study |
| pUA35–OSH6 | *OSH6* driven by the *TDH3* promoter, *URA3 CEN4* | This study |
| pUA35–OSH7 | *OSH7* driven by the *TDH3* promoter, *URA3 CEN4* | This study |
| YEplac195–OLE1 | *OLE1* driven by its own promoter, *URA3* 2μ | This study |

### 4.3. Confocal fluorescence microscopy

Cells expressing–mNG- or–mCherry-tagged proteins were visualized using a confocal laser-scanning microscope (FV-3000; Olympus Corporation, Tokyo, Japan) equipped with a UPLAPO60XOHR oil-immersion objective lens (numerical aperture = 1.5).

### 4.4. Lipid preparation after cultivation at high pressure and low temperature

Cells were inoculated at an initial OD_600_ adjusted so that, after 16 h of cultivation, all cultures reached the logarithmic growth phase at an OD_600_ of approximately 1.0. Cultures (15 mL) were placed in sterilized centrifuge tubes and sealed with Parafilm. Cells were grown under three conditions: 0.1 MPa at 25 °C, 25 MPa at 25 °C, and 0.1 MPa at 15 °C. After cultivation, cells were collected by centrifugation, washed twice with ice-cold phosphate-buffered saline (PBS) (FUJIFILM Wako Pure Chemical Corporation, Osaka, Japan), and subjected to lipidomic analysis. For ER membrane analysis, cells were first fractionated to obtain the S9 fraction, which was then centrifuged at 13,000 × *g* for 10 min to remove PM and mitochondria that sediment into the P13 pellet. Although the P13 fraction also contains ER membranes, it was discarded to avoid contamination, and the resulting supernatant was subsequently centrifuged at 100,000 × *g* for 1 h to obtain the P100 pellet, used as an ER-enriched microsomal fraction. This procedure followed previous reports. Zinser et al. demonstrated that pellets obtained at 20,000–30,000 × *g* contain PM and mitochondria together with ER membranes marked by GDP-mannosyltransferase (heavy microsomal fraction), whereas centrifugation at 100,000 × *g* for 1 h yields an ER fraction enriched in NADPH-cytochrome *c* reductase (light microsomal fraction). The vacuolar membrane marker α-D-mannosidase was absent from both ER fractions (Zinser *et al*., 1991). Reinhard et al. (2024) further developed the MemPrep method for high-purity organelle membrane isolation, using the P100 fraction as the starting material to minimize mitochondrial contamination (Reinhard *et al*., 2024). Based on these studies, the P100 fraction used in this work was considered to consist primarily of ER membranes, with minor contributions from Golgi and endosomal membranes. The P100 membrane fractions were subjected to lipidomic analysis.

Lipidomic analysis was performed using the *Lipidome Lab non-targeted Lipidome Scan* package (Lipidome Lab, Akita, Japan), employing liquid-chromatography (LC)–Orbitrap mass spectrometry (MS) as described previously (Nishiumi *et al*., 2022; Takumi *et al*., 2022). Internal standards were obtained from Avanti Polar Lipids (Alabaster, AL, USA). Methanol, isopropanol, and chloroform (Ultra–Performance LC/MS grade) were purchased from FUJIFILM Wako Pure Chemical Corporation, and ultrapure water was prepared using a Milli-Q system (Millipore, Milford, MA, USA). Briefly, cell pellets were suspended in methanol and homogenized. Lipids were extracted using a liquid–liquid extraction procedure based on the Bligh and Dyer method. The organic (lower) phase was evaporated under nitrogen, and the dried residues were re-dissolved in methanol for LC-MS/MS analysis.

### 4.5. Instrumental analysis and data processing

LC–electrospray ionization–MS/MS analysis was performed on a Q Exactive Plus mass spectrometer coupled to an UltiMate 3000 LC system (Thermo Fisher Scientific, Waltham, MA, USA). Separation was achieved on an L-column^3^ C18 metal–free column (2.0 µm, 2.0 mm × 100 mm i.d.) at 40 °C using the following solvent system: mobile phase A (isopropanol/methanol/water, 5:1:4 [v/v/v], containing 5 mM ammonium formate and 0.05% ammonium hydroxide [28% in water]) and mobile phase B (isopropanol containing 5 mM ammonium formate and 0.05% ammonium hydroxide [28% in water]). The gradient program was as follows: 60/40 (0 min), 40/60 (0–1 min), 20/80 (1–9 min), 5/95 (9–11 min), 5/95 (11–22 min), 95/5 (22–22.1 min), 95/5 (22.1–25 min), 60/40 (25–25.1 min), and 60/40 (25.1–30 min). The injection volume was 10 µL, and the flow rate was 0.1 mL/min.

Mass spectrometry was performed using a heated electrospray ionization (HESI-II) source under the following conditions: positive or negative ion mode; sheath gas, 60 arbitrary units; auxiliary gas, 10 arbitrary units; sweep gas, 0 arbitrary units; spray voltage, +3.2 kV (positive) or −3.0 kV (negative); heater temperature, 325 °C; ion transfer capillary temperature, 300 °C (positive) or 320 °C (negative); and S-lens RF level, 50. The Orbitrap analyzer operated at a resolution of 70,000 in full-scan mode (m/z 200–1,800; AGC target 1 × 10⁶ in positive and 3 × 10⁶ in negative mode) and at 17,500 (positive) or 35,000 (negative) in Top20 data-dependent MS^2^ mode (stepped normalized collision energy = 20, 30, and 40; isolation window = 4.0 m/z; AGC target = 1 × 10⁵; dynamic exclusion = 10 s).

Raw data were processed using LipidSearch 5.1.6 (Mitsui Knowledge Industries, Tokyo, Japan), which identifies intact lipid molecules based on molecular mass and MS/MS fragmentation of headgroups and fatty-acid chains. As appropriate internal standards were not available for all detected lipid species, biological matrix effects could not be normalized for every peak. Relative abundance values were calculated as the ratio of each analyte’s peak area to the total peak area. For selected lipid classes, normalization with internal standards was applied. Quantification and annotation followed the “absolute quantification Level 2 or 4” and “Fatty Acyl/Alkyl Level or Hydroxyl Group Level” schemes defined by the Lipidomics Standards Initiative (Lipidomics Standards Initiative Consortium, 2019).

### 4.6. Western blotting

Western blotting was performed for the whole cell extracts, P13 and P100 membrane fractions, as previously described, using an anti-HA monoclonal antibody (Medical and Biological Laboratories Co., Ltd., Nagoya, Japan), anti-Pma1 polyclonal antibody (Kato *et al*., 2025), anti-Pep2 monoclonal antibody (2C3G4, ab113689, Abcam Inc., Waltham MA, USA), anti-dolichol phosphate mannose synthase (Dpm1) monoclonal antibody (5C5, A6429, ThermoFisher Scientific), and horseradish peroxidase-conjugated anti-mouse IgG antibody (GE Healthcare Life Sciences, Piscataway, NJ, USA). Chemiluminescent signals were visualized using ImageQuant LAS4000 mini (GE Healthcare, Chicago, IL, USA).

### 4.7. Estimation of membrane curvature (PLCI) and lipid unsaturation index (DBI)

The phospholipid curvature index (PLCI), recently proposed by Winnikoff and colleagues (2024, 2025), was calculated for each yeast strain under the respective culture conditions. The lipid intrinsic curvature (*c*_0_) is a parameter representing the natural curvature adopted by a lipid monolayer in an unstressed state. To simplify curvature-based lipid composition analysis, these authors introduced the PLCI as an index derived from the weighted average of the intrinsic curvature values (*c*_0_,*i*) of individual lipid species, multiplied by their relative molar fractions (*x_i_*) (Winnikoff *et al*., 2024, Winnikoff and Budin, 2025).

In this study, reference *c*_0_ values were adopted from (Dymond, 2021) and (Mulet *et al*., 2008). Specifically, the following representative mean c_0_ values were used: PC = –0.01, PE = –0.035, PS = 0.005, and PI = –0.015 (in units of 1 Å^−1^). Using the lipidomic profiles of total cellular lipids and the P100 membrane fraction, the PLCI was calculated according to the following equation:

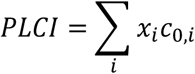

where *x_i_* is the molar fraction (mol%) of lipid species *i*, and *c_0,i_* is the intrinsic monolayer spontaneous curvature (1 Å^−1^) of lipid *i*. The PLCI represents the average intrinsic curvature of all phospholipid species in a membrane, weighted by their relative abundance.

The DBI, which indicates the overall degree of lipid unsaturation, was calculated using the following equation:

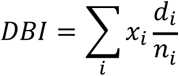

where *x_i_* is the molar fraction (mol%) of lipid species *i*, *d_i_* is the total number of double bonds per molecule, and *n_i_* is the number of acyl chains in that lipid.

### 4.8. Membrane physical property analysis after cultivation at high pressure and low temperature

Time-resolved fluorescence anisotropy of TMA–DPH was measured using a DeltaFlex spectrofluorometer (Horiba Scientific, Kyoto, Japan) equipped with a polarizing device and a 378 nm laser diode (NanoLED 375L, Horiba Scientific) operated at a pulse frequency of 1 MHz. The system was capable of time-correlated single-photon counting (TCSPC). Measurements were performed essentially as described previously using a FluoroCube (Horiba Scientific) (Abe and Hiraki, 2009; Abe *et al*., 2009), with the following modifications. In our earlier work, Tris buffer was used for anisotropy measurements; however, we found that Tris exhibits weak intrinsic fluorescence (excitation at 378 nm, emission at approximately 460 nm), which was negligible in steady-state experiments but detectable in TCSPC as a short-lifetime component, thereby interfering with the analysis. To eliminate this artifact, PBS buffer was used, which showed no detectable fluorescence under the same conditions and substantially improved the reliability of lifetime measurements.

For membrane fluidity measurements, cells grown in SC medium under each condition were washed twice with cold Milli-Q water, resuspended in PBS containing 1 mM EDTA (pH 7.3), and labeled with 0.5 µM TMA-DPH for 5 min at 25 °C in the dark, allowing the probe to partition into the outer leaflet region of the PM lipid bilayer. After labeling, the cells were washed twice with cold Milli-Q water, resuspended in PBS/1 mM EDTA, and kept on ice until measurement. For P100 membrane fractions, 10⁸ cells were disrupted with 0.3 g glass beads (425–600 µm) by vortexing at 4 °C for 5 min in PBS/1 mM EDTA. The supernatant was labeled with TMA-DPH (final 0.5 µM) at 25 °C in the dark, then centrifuged at 13,000 × *g* for 10 min (to remove the PM, vacuolar membrane, and mitochondria), and subsequently at 100,000 × *g* for 1 h to obtain the P100 membrane pellet. The pellet was used as the TMA-DPH–labeled P100 membrane sample. Cell and membrane suspensions were diluted in PBS/1 mM EDTA to maintain photon counts below 20,000 photons s^−1^, and the measurements were performed at 25 °C. The time axis ranged from 0 to 200 ns with 4,000 bins (50 ps/bin). *I*_VH(*t*)_ and *I*_VV(*t*)_ were sequentially measured, and the G-factor was determined from *I*_HH(*t*)_ and *I*_HV(*t*)_ recorded for 2 min each. The instrumental response function was obtained by measuring Rayleigh scattering from a colloidal silica suspension at 378 nm. Decay parameters for fluorescence intensity and anisotropy were determined by convolution of *I*_VH(*t*)_ and *I*_VV(*t*)_ with the instrument response function. Data analysis was performed using EzTime software version 3.4.1.1 (Horiba Scientific, Kyoto, Japan).

Anisotropy decay, *r* (*t*), was approximated using a single-exponential model of fluorophore motion restricted in a cone:

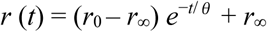

where *r_0_* is the initial anisotropy, *r*_∞_ the limiting anisotropy, and *θ* (ns) the rotational correlation time. The order parameter *S*, providing structural information about the membrane, i.e., the mean orientational order of the lipid chains, was calculated as:

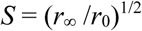

Fluorescence decay, *I* (*t*), was fitted to a three-exponential function:

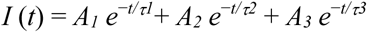

where *τ_i_* (ns) is the lifetime and *A_i_* the amplitude of each component. The amplitude-weighted mean fluorescence lifetime (< τ >) was calculated as:

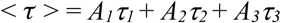

## Acknowledgements

We thank Dr. Roland Winter for critical reading of the manuscript and for valuable comments. We are also grateful to Drs. Hitoshi Matsuki, Kuninori Suzuki, Satoshi Uemura, Yoichi Noda, Ryoichi Fukuda, and Satoru Okada for their helpful discussions, and to Ayane Takemoto and other members of our laboratory for their cooperation throughout this study. Lipidomic analyses were conducted by Dr. Hiroki Nakanishi (Lipidome Lab) on a commissioned basis. We also thank Editage (www.editage.jp) for English language editing.

## Author’s contribution

Fumiyoshi Abe: Conceptualization, investigation, and manuscript writing. Tetsuo Mioka: Conceptualization and investigation.

Yusuke Suzuki, Akane Ogasawara, Daisuke Mochizuki, Saki Imura, Yusuke Kato, and Takahiro Mochizuki: Investigation.

## Data availability

All data supporting the findings of this study are included in the article and are available from the corresponding author upon reasonable request.

## Abbreviations

MPa: megapascals
BLTP: Bridge-like Lipid Transfer Protein
PLCI: phospholipid curvature index
DBI: double bond index
*S*: order parameter
<*τ*>: mean fluorescence lifetime of TMA-DPH

---

**Fig. EV1.**
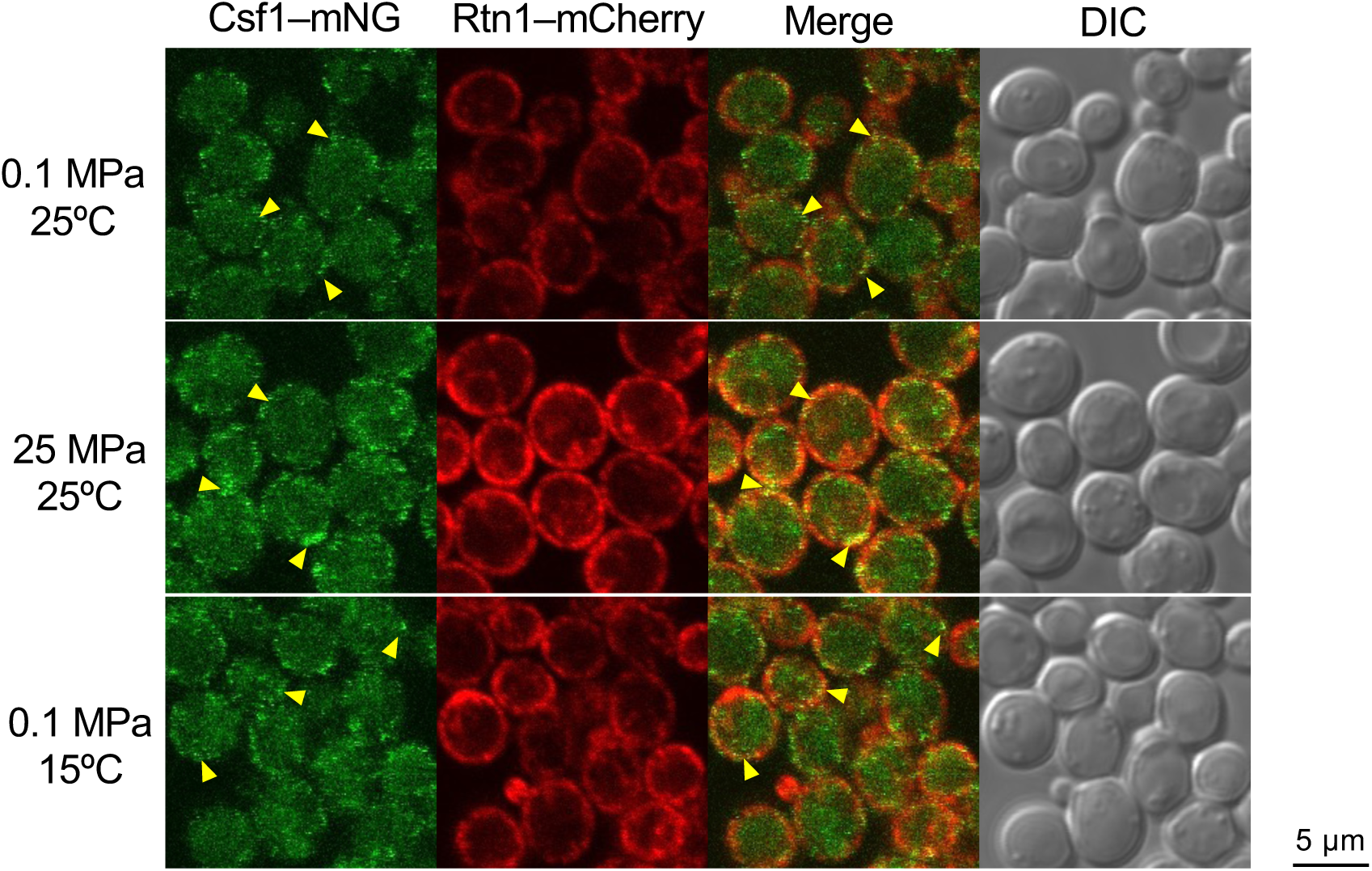
Colocalization of Csf1 with the cortical ER marker Rtn1. Cells co-expressing Csf1–mNG and Rtn1–mCherry were grown under the indicated conditions for 6 h and observed by confocal microscopy under atmospheric pressure. Yellow arrowheads indicate colocalization of the two proteins.

**Fig. EV2.**
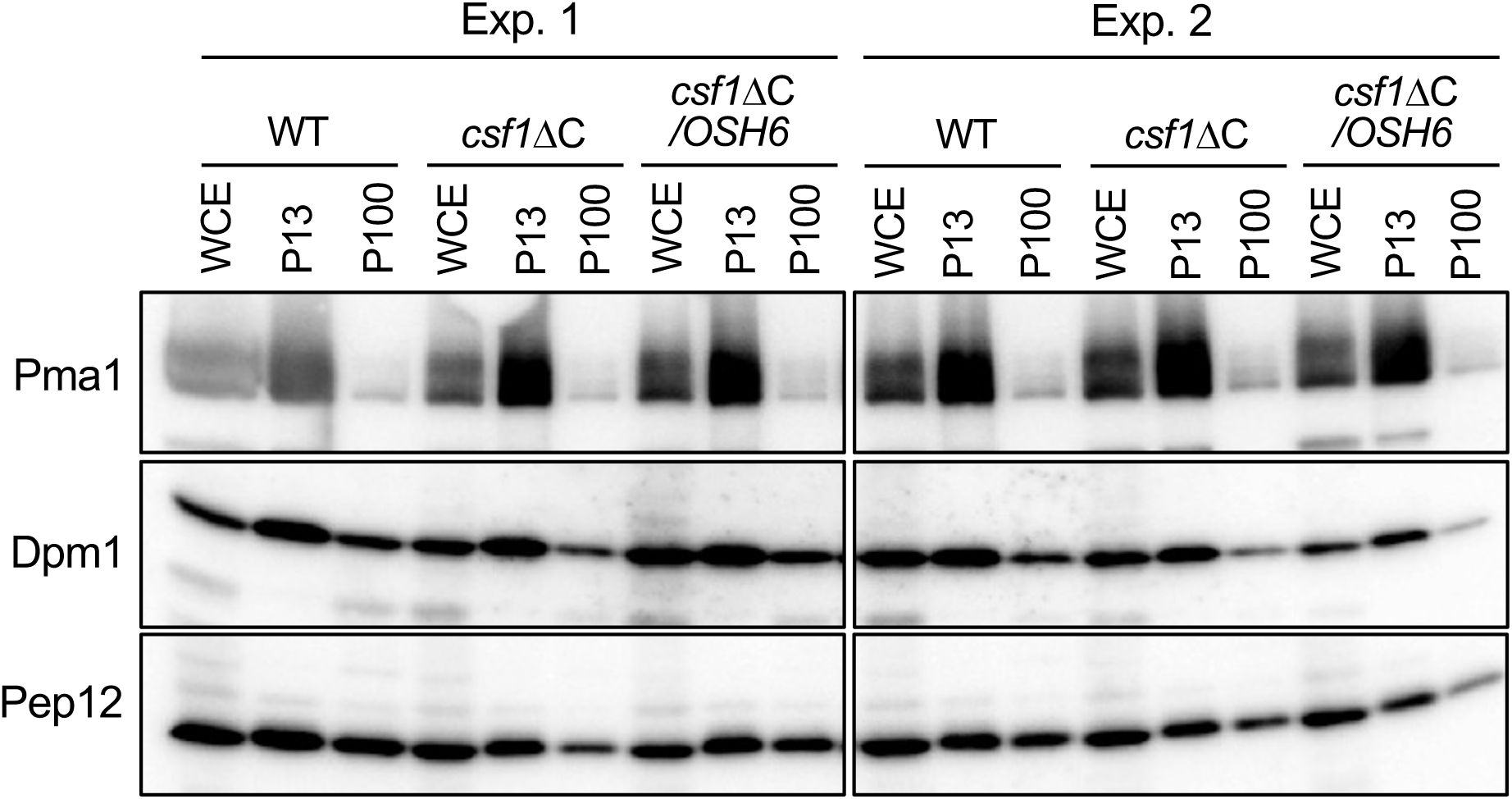
Detection of marker proteins in each membrane fraction. Whole-cell extracts, P13 and P100 membrane fractions prepared from cells were analyzed by Western blotting to detect marker proteins (Pma1 for the PM, Dpm1 for the ER membrane, and Pep12 for endosomes). WCE, whole cell extracts.

**Fig. EV3.**
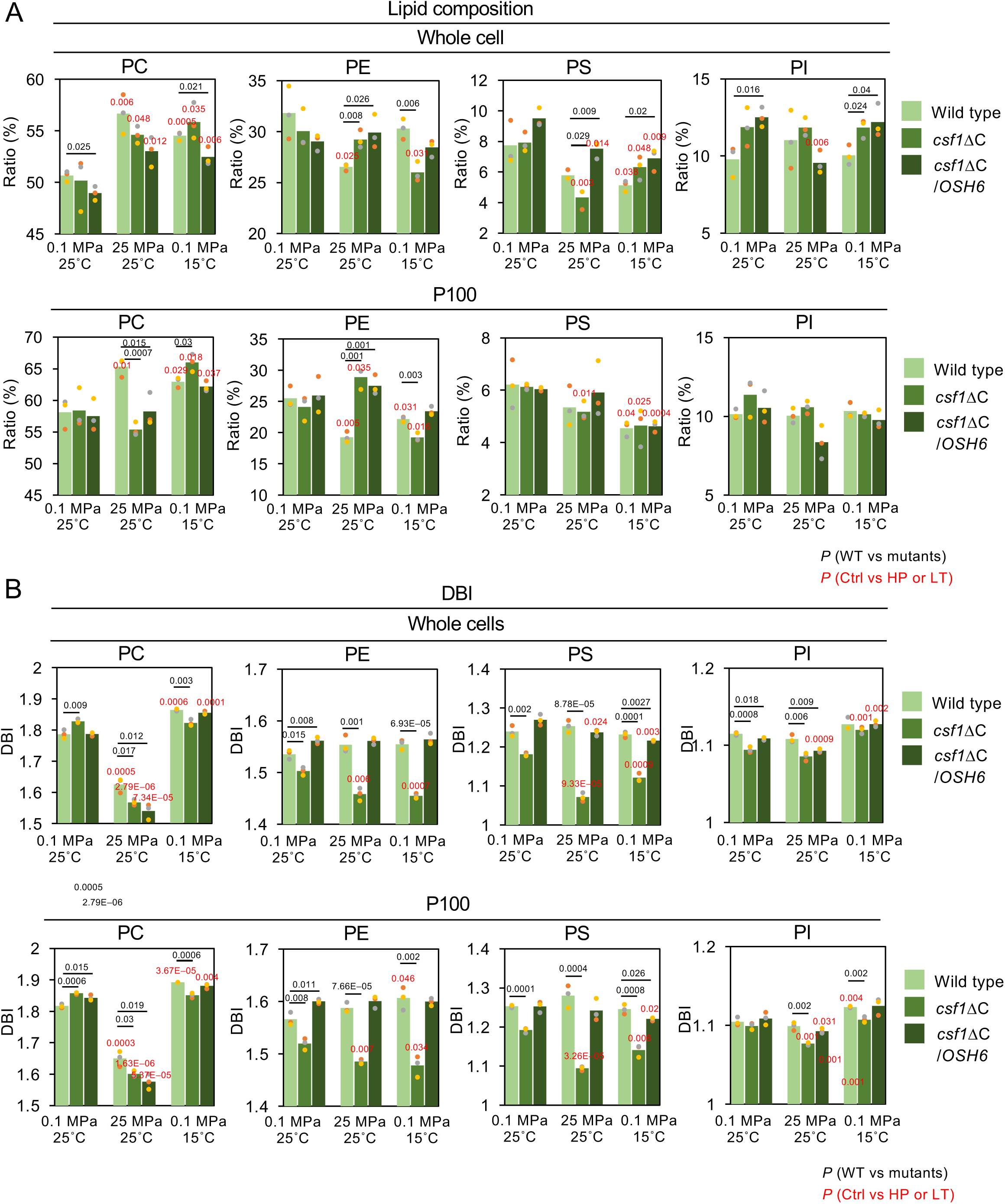
Lipid composition and DBI of cells and P100 membrane fractions. Changes in phospholipid composition (A) and DBI (B) after 16 h of growth to OD_600_ around 1.0 under indicated conditions. Phospholipids were quantified by lipidomic analysis. Data represent relative amounts of phospholipids containing C16 and C18 acyl chains from three independent preparations. PC, phosphatidylcholine; PE, phosphatidylethanolamine; PI, phosphatidylinositol; PS, phosphatidylserine. Absolute values of the bar graphs are provided in Table EV1 to EV4.

**Fig. EV4.**
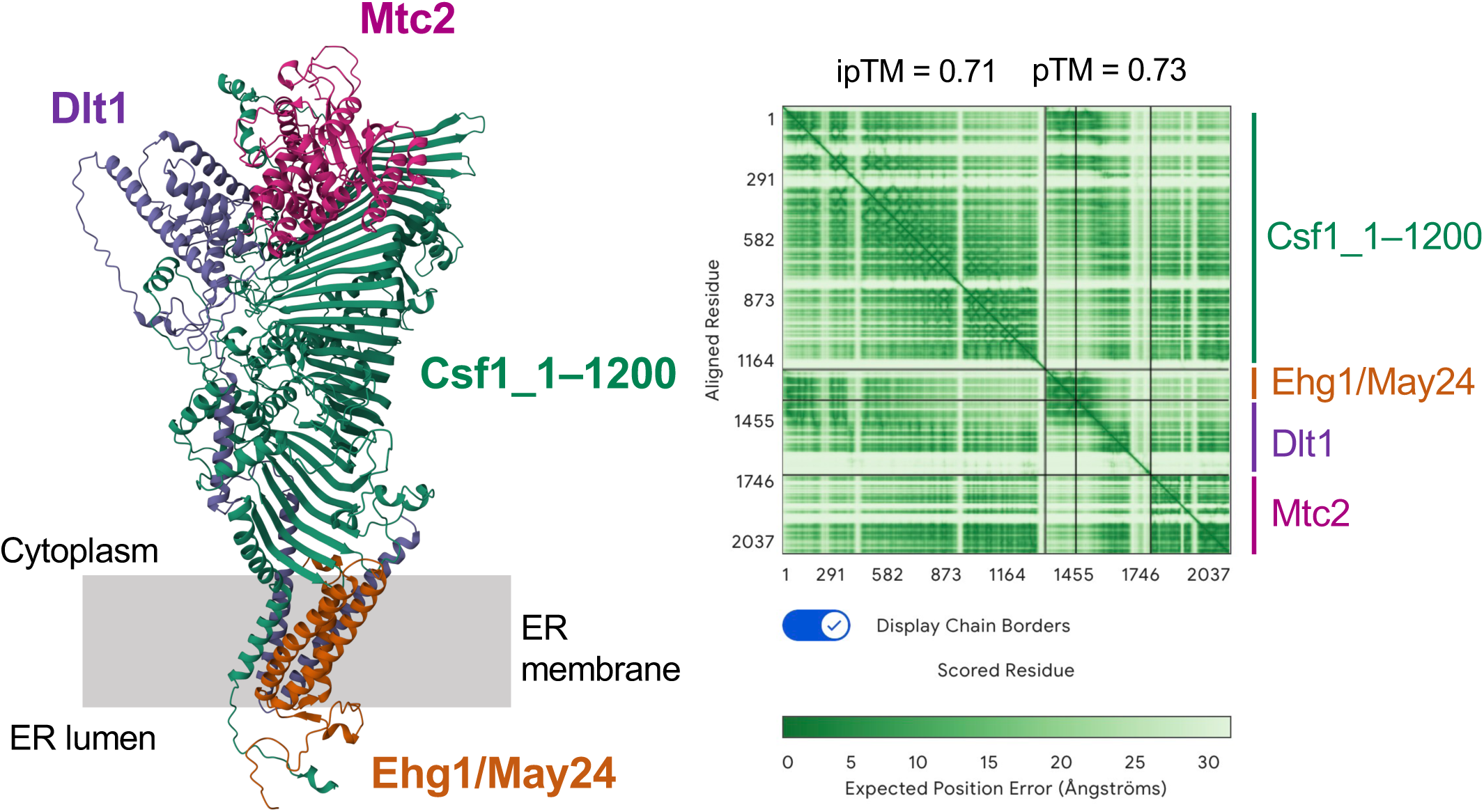
AlphaFold3-predicted complex of Csf1 (1–1200 aa), Ehg1/May24, Dlt1, and Mtc2. AlphaFold3 prediction of a single multiprotein complex composed of Csf1 (residues 1–1200), Ehg1/May24, Dlt1, and Mtc2. The left panel shows the integrated complex visualized with Mol* Viewer, illustrating the spatial arrangement of all four proteins. The right panel presents the expected position error map corresponding to the prediction confidence. The ipTM and pTM scores indicate the predicted interface confidence and overall structural accuracy, respectively. Importantly, deletion of any one of these four proteins, Csf1, Ehg1, Dlt1, or Mtc2, results in similar high-pressure and low-temperature sensitivity, supporting the possibility that they function cooperatively as a single complex to maintain membrane homeostasis under stress.

**Table EV1.** Composition of whole-cell lipids.

| Lipid sp. | Condition | Strain | Mean (mol %) | No. 1 (mol %) | No. 2 (mol %) | No. 3 (mol %) | P (Mutant vs WT) | P (HP, LT vs Ctrl) |
| --- | --- | --- | --- | --- | --- | --- | --- | --- |
| PC | 0.1 MPa, 25°C | Wild | 50.7 | 50.8 | 51.0 | 50.1 |  |  |
| PC | 0.1 MPa, 25°C | *csf1ΔC* | 50.2 | 51.8 | 51.5 | 47.2 | 0.76284 |  |
| PC | 0.1 MPa, 25°C | *csf1ΔC/OSH6* | 49.0 | 49.0 | 49.6 | 48.3 | 0.02484 |  |
| PC | 25 MPa, 25°C | Wild | 56.7 | 58.5 | 56.8 | 54.7 |  | 0.00614 |
| PC | 25 MPa, 25°C | *csf1ΔC* | 54.6 | 54.6 | 55.5 | 53.8 | 0.16109 | 0.04771 |
| PC | 25 MPa, 25°C | *csf1ΔC/OSH6* | 53.0 | 53.2 | 54.4 | 51.5 | 0.05746 | 0.01156 |
| PC | 0.1 MPa, 15°C | Wild | 54.5 | 54.8 | 54.7 | 54.1 |  | 0.00047 |
| PC | 0.1 MPa, 15°C | *csf1ΔC* | 55.8 | 55.5 | 57.7 | 54.3 | 0.27093 | 0.03470 |
| PC | 0.1 MPa, 15°C | *csf1ΔC/OSH6* | 52.5 | 53.5 | 51.8 | 52.1 | 0.02110 | 0.00551 |
| PE | 0.1 MPa, 25°C | Wild | 31.8 | 29.3 | 31.7 | 34.5 |  |  |
| PE | 0.1 MPa, 25°C | *csf1ΔC* | 30.0 | 28.9 | 29.0 | 32.3 | 0.39550 |  |
| PE | 0.1 MPa, 25°C | *csf1ΔC/OSH6* | 29.0 | 29.4 | 28.1 | 29.6 | 0.15037 |  |
| PE | 25 MPa, 25°C | Wild | 26.5 | 26.1 | 26.7 | 26.7 |  | 0.02494 |
| PE | 25 MPa, 25°C | *csf1ΔC* | 29.2 | 30.2 | 28.5 | 29.0 | 0.00772 | 0.53544 |
| PE | 25 MPa, 25°C | *csf1ΔC/OSH6* | 29.9 | 28.6 | 29.4 | 31.7 | 0.02551 | 0.45663 |
| PE | 0.1 MPa, 15°C | Wild | 30.3 | 29.3 | 30.4 | 31.3 |  | 0.39785 |
| PE | 0.1 MPa, 15°C | *csf1ΔC* | 26.0 | 25.3 | 25.6 | 27.1 | 0.00573 | 0.03122 |
| PE | 0.1 MPa, 15°C | *csf1ΔC/OSH6* | 28.4 | 29.1 | 27.5 | 28.8 | 0.06799 | 0.42680 |
| PS | 0.1 MPa, 25°C | Wild | 7.7 | 9.4 | 7.0 | 6.8 |  |  |
| PS | 0.1 MPa, 25°C | *csf1ΔC* | 7.9 | 8.6 | 7.7 | 7.4 | 0.85070 |  |
| PS | 0.1 MPa, 25°C | *csf1ΔC/OSH6* | 9.5 | 9.2 | 9.1 | 10.2 | 0.12256 |  |
| PS | 25 MPa, 25°C | Wild | 5.8 | 6.2 | 5.6 | 5.6 |  | 0.08333 |
| PS | 25 MPa, 25°C | *csf1ΔC* | 4.3 | 3.6 | 4.7 | 4.7 | 0.02925 | 0.00253 |
| PS | 25 MPa, 25°C | *csf1ΔC/OSH6* | 7.5 | 7.9 | 6.9 | 7.8 | 0.00912 | 0.01375 |
| PS | 0.1 MPa, 15°C | Wild | 5.1 | 5.3 | 5.4 | 4.7 |  | 0.03848 |
| PS | 0.1 MPa, 15°C | *csf1ΔC* | 6.3 | 7.0 | 5.5 | 6.5 | 0.07104 | 0.04817 |
| PS | 0.1 MPa, 15°C | *csf1ΔC/OSH6* | 6.9 | 6.1 | 7.2 | 7.4 | 0.02009 | 0.00893 |
| PI | 0.1 MPa, 25°C | Wild | 9.8 | 10.4 | 10.3 | 8.6 |  |  |
| PI | 0.1 MPa, 25°C | *csf1ΔC* | 11.9 | 10.6 | 11.8 | 13.1 | 0.08634 |  |
| PI | 0.1 MPa, 25°C | *csf1ΔC/OSH6* | 12.5 | 12.4 | 13.2 | 11.9 | 0.01616 |  |
| PI | 25 MPa, 25°C | Wild | 11.0 | 9.2 | 10.9 | 13.0 |  | 0.37154 |
| PI | 25 MPa, 25°C | *csf1ΔC* | 11.8 | 11.7 | 11.3 | 12.5 | 0.51438 | 0.97734 |
| PI | 25 MPa, 25°C | *csf1ΔC/OSH6* | 9.6 | 10.4 | 9.3 | 9.0 | 0.27881 | 0.00624 |
| PI | 0.1 MPa, 15°C | Wild | 10.0 | 10.7 | 9.5 | 10.0 |  | 0.71935 |
| PI | 0.1 MPa, 15°C | *csf1ΔC* | 11.8 | 12.2 | 11.1 | 12.1 | 0.02420 | 0.97406 |
| PI | 0.1 MPa, 15°C | *csf1ΔC/OSH6* | 12.2 | 11.4 | 13.4 | 11.8 | 0.03964 | 0.69028 |

Cells were cultured under each condition for 16 h.
Ctrl, 0.1 MPa, 25°C; HP, 0.1 MPa, 25°C, LT, 0.1 MPa, 15°C
P, P value of Student's *t*-test

**Table EV2.** Composition of P100-membrane lipids.

| Lipid sp. | Condition | Strain | Mean (mol %) | No. 1 (mol %) | No. 2 (mol %) | No. 3 (mol %) | P (Mutant vs WT) | P (HP, LT vs Ctrl) |
| --- | --- | --- | --- | --- | --- | --- | --- | --- |
| PC | 0.1 MPa, 25°C | Wild | 58.1 | 55.4 | 59.7 | 59.2 |  |  |
| PC | 0.1 MPa, 25°C | *csf1ΔC* | 58.4 | 56.4 | 56.8 | 62.0 | 0.91106 |  |
| PC | 0.1 MPa, 25°C | *csf1ΔC/OSH6* | 57.5 | 55.4 | 56.8 | 60.3 | 0.76854 |  |
| PC | 25 MPa, 25°C | Wild | 65.3 | 63.6 | 66.1 | 66.3 |  | 0.01075 |
| PC | 25 MPa, 25°C | *csf1ΔC* | 55.4 | 55.0 | 54.6 | 56.6 | 0.00066 | 0.18697 |
| PC | 25 MPa, 25°C | *csf1ΔC/OSH6* | 58.2 | 56.9 | 61.3 | 56.5 | 0.01527 | 0.74206 |
| PC | 0.1 MPa, 15°C | Wild | 62.9 | 62.0 | 63.3 | 63.5 |  | 0.02899 |
| PC | 0.1 MPa, 15°C | *csf1ΔC* | 66.0 | 66.1 | 67.3 | 64.6 | 0.02978 | 0.01807 |
| PC | 0.1 MPa, 15°C | *csf1ΔC/OSH6* | 62.2 | 63.1 | 61.5 | 61.9 | 0.33457 | 0.03743 |
| PE | 0.1 MPa, 25°C | Wild | 25.5 | 27.5 | 24.5 | 24.6 |  |  |
| PE | 0.1 MPa, 25°C | *csf1ΔC* | 24.1 | 25.5 | 25.0 | 21.8 | 0.40425 |  |
| PE | 0.1 MPa, 25°C | *csf1ΔC/OSH6* | 25.9 | 29.0 | 25.5 | 23.3 | 0.84503 |  |
| PE | 25 MPa, 25°C | Wild | 19.3 | 20.2 | 19.1 | 18.5 |  | 0.00471 |
| PE | 25 MPa, 25°C | *csf1ΔC* | 28.9 | 29.9 | 29.8 | 26.9 | 0.00095 | 0.03480 |
| PE | 25 MPa, 25°C | *csf1ΔC/OSH6* | 27.5 | 29.2 | 26.3 | 26.9 | 0.00126 | 0.45195 |
| PE | 0.1 MPa, 15°C | Wild | 22.2 | 22.5 | 22.4 | 21.7 |  | 0.03061 |
| PE | 0.1 MPa, 15°C | *csf1ΔC* | 19.2 | 18.8 | 18.9 | 20.0 | 0.00296 | 0.01570 |
| PE | 0.1 MPa, 15°C | *csf1ΔC/OSH6* | 23.4 | 23.1 | 24.2 | 22.9 | 0.06022 | 0.21623 |
| PS | 0.1 MPa, 25°C | Wild | 6.2 | 7.2 | 5.3 | 6.2 |  |  |
| PS | 0.1 MPa, 25°C | *csf1ΔC* | 6.1 | 6.1 | 6.0 | 6.2 | 0.87783 |  |
| PS | 0.1 MPa, 25°C | *csf1ΔC/OSH6* | 6.0 | 6.0 | 6.1 | 6.1 | 0.76001 |  |
| PS | 25 MPa, 25°C | Wild | 5.3 | 6.2 | 5.2 | 4.7 |  | 0.27501 |
| PS | 25 MPa, 25°C | *csf1ΔC* | 5.2 | 4.9 | 5.0 | 5.6 | 0.74452 | 0.01070 |
| PS | 25 MPa, 25°C | *csf1ΔC/OSH6* | 5.9 | 5.5 | 5.1 | 7.1 | 0.49771 | 0.84408 |
| PS | 0.1 MPa, 15°C | Wild | 4.5 | 4.7 | 4.2 | 4.7 |  | 0.03976 |
| PS | 0.1 MPa, 15°C | *csf1ΔC* | 4.7 | 4.9 | 3.8 | 5.2 | 0.83051 | 0.02497 |
| PS | 0.1 MPa, 15°C | *csf1ΔC/OSH6* | 4.6 | 4.4 | 4.7 | 4.8 | 0.73614 | 0.00035 |
| PI | 0.1 MPa, 25°C | Wild | 10.1 | 9.9 | 10.5 | 10.0 |  |  |
| PI | 0.1 MPa, 25°C | *csf1ΔC* | 11.4 | 12.0 | 12.1 | 10.0 | 0.16616 |  |
| PI | 0.1 MPa, 25°C | *csf1ΔC/OSH6* | 10.5 | 9.6 | 11.6 | 10.3 | 0.54755 |  |
| PI | 25 MPa, 25°C | Wild | 10.0 | 10.0 | 9.6 | 10.5 |  | 0.78385 |
| PI | 25 MPa, 25°C | *csf1ΔC* | 10.6 | 10.2 | 10.6 | 11.0 | 0.20799 | 0.34668 |
| PI | 25 MPa, 25°C | *csf1ΔC/OSH6* | 8.4 | 8.3 | 7.3 | 9.4 | 0.06561 | 0.06128 |
| PI | 0.1 MPa, 15°C | Wild | 10.3 | 10.9 | 10.1 | 10.0 |  | 0.57192 |
| PI | 0.1 MPa, 15°C | *csf1ΔC* | 10.1 | 10.2 | 10.0 | 10.2 | 0.50698 | 0.15637 |
| PI | 0.1 MPa, 15°C | *csf1ΔC/OSH6* | 9.8 | 9.3 | 9.5 | 10.4 | 0.26105 | 0.31741 |

Cells were cultured under each condition for 16 h.
Ctrl, 0.1 MPa, 25°C; HP, 0.1 MPa, 25°C, LT, 0.1 MPa, 15°C
P, P value of Student's *t* test

**Table EV3.** DBI values of whole-cell lipids.

| Lipid sp. | Condition | Strain | Mean | No. 1 | No. 2 | No. 3 | P (Mutant vs WT) | P (HP, LT vs Ctrl) |
| --- | --- | --- | --- | --- | --- | --- | --- | --- |
| PC | 0.1 MPa, 25°C | Wild | 1.786 | 1.785 | 1.800 | 1.773 |  |  |
| PC | 0.1 MPa, 25°C | *csf1ΔC* | 1.827 | 1.822 | 1.836 | 1.824 | 0.00928 |  |
| PC | 0.1 MPa, 25°C | *csf1ΔC/OSH6* | 1.787 | 1.780 | 1.790 | 1.791 | 0.92560 |  |
| PC | 25 MPa, 25°C | Wild | 1.625 | 1.598 | 1.638 | 1.639 |  | 0.00049 |
| PC | 25 MPa, 25°C | *csf1ΔC* | 1.567 | 1.558 | 1.577 | 1.567 | 0.01659 | 0.00000 |
| PC | 25 MPa, 25°C | *csf1ΔC/OSH6* | 1.540 | 1.558 | 1.549 | 1.512 | 0.01217 | 0.00007 |
| PC | 0.1 MPa, 15°C | Wild | 1.864 | 1.864 | 1.861 | 1.867 |  | 0.00061 |
| PC | 0.1 MPa, 15°C | *csf1ΔC* | 1.822 | 1.819 | 1.834 | 1.814 | 0.00262 | 0.51728 |
| PC | 0.1 MPa, 15°C | *csf1ΔC/OSH6* | 1.855 | 1.860 | 1.854 | 1.850 | 0.05812 | 0.00015 |
| PE | 0.1 MPa, 25°C | Wild | 1.536 | 1.539 | 1.526 | 1.542 |  |  |
| PE | 0.1 MPa, 25°C | *csf1ΔC* | 1.503 | 1.505 | 1.510 | 1.495 | 0.00831 |  |
| PE | 0.1 MPa, 25°C | *csf1ΔC/OSH6* | 1.562 | 1.556 | 1.561 | 1.568 | 0.01486 |  |
| PE | 25 MPa, 25°C | Wild | 1.554 | 1.572 | 1.542 | 1.547 |  | 0.17258 |
| PE | 25 MPa, 25°C | *csf1ΔC* | 1.458 | 1.459 | 1.470 | 1.445 | 0.00128 | 0.00592 |
| PE | 25 MPa, 25°C | *csf1ΔC/OSH6* | 1.561 | 1.563 | 1.554 | 1.567 | 0.50537 | 0.94024 |
| PE | 0.1 MPa, 15°C | Wild | 1.555 | 1.562 | 1.559 | 1.545 |  | 0.06030 |
| PE | 0.1 MPa, 15°C | *csf1ΔC* | 1.455 | 1.459 | 1.455 | 1.451 | 0.00007 | 0.00071 |
| PE | 0.1 MPa, 15°C | *csf1ΔC/OSH6* | 1.564 | 1.560 | 1.576 | 1.556 | 0.32707 | 0.73988 |
| PS | 0.1 MPa, 25°C | Wild | 1.240 | 1.255 | 1.235 | 1.228 |  |  |
| PS | 0.1 MPa, 25°C | *csfI*ΔC | 1.181 | 1.186 | 1.181 | 1.177 | 0.00231 |  |
| PS | 0.1 MPa, 25°C | *csfI*ΔC/*OSH6* | 1.270 | 1.256 | 1.284 | 1.268 | 0.05870 |  |
| PS | 25 MPa, 25°C | Wild | 1.254 | 1.269 | 1.255 | 1.237 |  | 0.32160 |
| PS | 25 MPa, 25°C | *csfI*ΔC | 1.072 | 1.061 | 1.084 | 1.069 | 0.00009 | 0.00009 |
| PS | 25 MPa, 25°C | *csfI*ΔC/*OSH6* | 1.237 | 1.243 | 1.229 | 1.240 | 0.18808 | 0.02413 |
| PS | 0.1 MPa, 15°C | Wild | 1.232 | 1.224 | 1.239 | 1.234 |  | 0.46643 |
| PS | 0.1 MPa, 15°C | *csfI*ΔC | 1.121 | 1.134 | 1.116 | 1.114 | 0.00013 | 0.00085 |
| PS | 0.1 MPa, 15°C | *csfI*ΔC/*OSH6* | 1.216 | 1.214 | 1.217 | 1.218 | 0.02700 | 0.00280 |
| PI | 0.1 MPa, 25°C | Wild | 1.115 | 1.116 | 1.113 | 1.115 |  |  |
| PI | 0.1 MPa, 25°C | *csfI*ΔC | 1.094 | 1.091 | 1.093 | 1.098 | 0.00084 |  |
| PI | 0.1 MPa, 25°C | *csfI*ΔC/*OSH6* | 1.109 | 1.110 | 1.107 | 1.110 | 0.01818 |  |
| PI | 25 MPa, 25°C | Wild | 1.108 | 1.114 | 1.104 | 1.106 |  | 0.10137 |
| PI | 25 MPa, 25°C | *csfI*ΔC | 1.086 | 1.080 | 1.087 | 1.090 | 0.00590 | 0.07745 |
| PI | 25 MPa, 25°C | *csfI*ΔC/*OSH6* | 1.093 | 1.091 | 1.092 | 1.096 | 0.00942 | 0.00089 |
| PI | 0.1 MPa, 15°C | Wild | 1.127 | 1.123 | 1.137 | 1.122 |  | 0.06199 |
| PI | 0.1 MPa, 15°C | *csfI*ΔC | 1.120 | 1.124 | 1.115 | 1.121 | 0.24369 | 0.00138 |
| PI | 0.1 MPa, 15°C | *csfI*ΔC/*OSH6* | 1.128 | 1.126 | 1.124 | 1.132 | 0.97152 | 0.00208 |

Cells were cultured under each condition for 16 h.
Ctrl, 0.1 MPa, 25°C; HP, 0.1 MPa, 25°C, LT, 0.1 MPa, 15°C
P, P value of Student's *t*-test

**Table EV4.** DBI values of P100-membrane lipids.

| Lipid sp. | Condition | Strain | No. 1 | No. 1 | No. 2 | No. 3 | P (Mutant vs WT) | P (HP, LT vs Ctrl) |
| --- | --- | --- | --- | --- | --- | --- | --- | --- |
| PC | 0.1 MPa, 25°C | Wild | 1.817 | 1.812 | 1.825 | 1.815 |  |  |
| PC | 0.1 MPa, 25°C | *csf1ΔC* | 1.858 | 1.854 | 1.860 | 1.859 | 0.00063 |  |
| PC | 0.1 MPa, 25°C | *csf1ΔC/OSH6* | 1.843 | 1.852 | 1.840 | 1.836 | 0.01470 |  |
| PC | 25 MPa, 25°C | Wild | 1.650 | 1.625 | 1.654 | 1.673 |  | 0.00032 |
| PC | 25 MPa, 25°C | *csf1ΔC* | 1.601 | 1.597 | 1.594 | 1.612 | 0.03012 | 0.00000 |
| PC | 25 MPa, 25°C | *csf1ΔC/OSH6* | 1.576 | 1.576 | 1.601 | 1.553 | 0.01932 | 0.00005 |
| PC | 0.1 MPa, 15°C | Wild | 1.892 | 1.893 | 1.892 | 1.892 |  | 0.00004 |
| PC | 0.1 MPa, 15°C | *csf1ΔC* | 1.851 | 1.843 | 1.853 | 1.858 | 0.00063 | 0.23311 |
| PC | 0.1 MPa, 15°C | *csf1ΔC/OSH6* | 1.881 | 1.872 | 1.886 | 1.884 | 0.05810 | 0.00433 |
| PE | 0.1 MPa, 25°C | Wild | 1.566 | 1.561 | 1.580 | 1.557 |  |  |
| PE | 0.1 MPa, 25°C | *csf1ΔC* | 1.520 | 1.529 | 1.522 | 1.508 | 0.00840 |  |
| PE | 0.1 MPa, 25°C | *csf1ΔC/OSH6* | 1.600 | 1.596 | 1.604 | 1.599 | 0.01065 |  |
| PE | 25 MPa, 25°C | Wild | 1.588 | 1.583 | 1.599 | 1.581 |  | 0.07626 |
| PE | 25 MPa, 25°C | *csf1ΔC* | 1.485 | 1.489 | 1.485 | 1.481 | 0.00008 | 0.00700 |
| PE | 25 MPa, 25°C | *csf1ΔC/OSH6* | 1.600 | 1.605 | 1.587 | 1.609 | 0.21571 | 0.95083 |
| PE | 0.1 MPa, 15°C | Wild | 1.607 | 1.627 | 1.584 | 1.609 |  | 0.04648 |
| PE | 0.1 MPa, 15°C | *csf1ΔC* | 1.478 | 1.494 | 1.483 | 1.456 | 0.00160 | 0.03376 |
| PE | 0.1 MPa, 15°C | *csf1ΔC/OSH6* | 1.600 | 1.601 | 1.606 | 1.592 | 0.61713 | 0.93408 |
| PS | 0.1 MPa, 25°C | Wild | 1.253 | 1.254 | 1.249 | 1.256 |  |  |
| PS | 0.1 MPa, 25°C | *csfI*ΔC | 1.192 | 1.197 | 1.185 | 1.195 | 0.00013 |  |
| PS | 0.1 MPa, 25°C | *csfI*ΔC/*OSH6* | 1.253 | 1.264 | 1.237 | 1.258 | 0.96039 |  |
| PS | 25 MPa, 25°C | Wild | 1.280 | 1.305 | 1.286 | 1.249 |  | 0.17620 |
| PS | 25 MPa, 25°C | *csfI*ΔC | 1.094 | 1.088 | 1.099 | 1.096 | 0.00037 | 0.00003 |
| PS | 25 MPa, 25°C | *csfI*ΔC/*OSH6* | 1.242 | 1.220 | 1.233 | 1.274 | 0.17114 | 0.58516 |
| PS | 0.1 MPa, 15°C | Wild | 1.246 | 1.234 | 1.245 | 1.258 |  | 0.36912 |
| PS | 0.1 MPa, 15°C | *csfI*ΔC | 1.141 | 1.149 | 1.151 | 1.123 | 0.00078 | 0.00628 |
| PS | 0.1 MPa, 15°C | *csfI*ΔC/*OSH6* | 1.221 | 1.217 | 1.222 | 1.224 | 0.02624 | 0.02002 |
| PI | 0.1 MPa, 25°C | Wild | 1.104 | 1.100 | 1.110 | 1.102 |  |  |
| PI | 0.1 MPa, 25°C | *csfI*ΔC | 1.099 | 1.102 | 1.101 | 1.094 | 0.25592 |  |
| PI | 0.1 MPa, 25°C | *csfI*ΔC/*OSH6* | 1.109 | 1.116 | 1.109 | 1.100 | 0.46707 |  |
| PI | 25 MPa, 25°C | Wild | 1.099 | 1.093 | 1.100 | 1.103 |  | 0.25094 |
| PI | 25 MPa, 25°C | *csfI*ΔC | 1.077 | 1.078 | 1.075 | 1.078 | 0.00186 | 0.00100 |
| PI | 25 MPa, 25°C | *csfI*ΔC/*OSH6* | 1.092 | 1.092 | 1.090 | 1.096 | 0.13204 | 0.03053 |
| PI | 0.1 MPa, 15°C | Wild | 1.123 | 1.123 | 1.122 | 1.124 |  | 0.00366 |
| PI | 0.1 MPa, 15°C | *csfI*ΔC | 1.107 | 1.107 | 1.111 | 1.104 | 0.00177 | 0.06300 |
| PI | 0.1 MPa, 15°C | *csfI*ΔC/*OSH6* | 1.125 | 1.112 | 1.132 | 1.129 | 0.80242 | 0.10820 |

Cells were cultured under each condition for 16 h.
Ctrl, 0.1 MPa, 25°C; HP, 0.1 MPa, 25°C, LT, 0.1 MPa, 15°C
P, P value of Student's *t*-test

